# Sunscreen redox status in a multicellular cyanobacterium visualized by Raman scattering spectral microscope

**DOI:** 10.1101/2022.07.26.501630

**Authors:** Kouto Tamamizu, Toshio Sakamoto, Yuki Kurashige, Shuho Nozue, Shigeichi Kumazaki

## Abstract

UV radiation, desiccation, and starvation induce some cyanobacteria to produce a UVA-absorbing pigment, scytonemin, at extracellular sheaths. Although the accumulation of scytonemin is recognizable as dark sheaths through optical microscopes, it has been nontrivial to identify its redox status and obtain its subcellular distribution in response to physiological conditions. Here, we show that a spontaneous Raman scattering spectral microscopy based on an excitation-laser-line-scanning method unveil 3D subcellular distributions of non-UV-induced scytonemins with distinct redox statuses in a filamentous cyanobacterium with a single nitrogen-fixing cell at the basal end. Cellular differentiations and scytonemin redox conditions were simultaneously visualized with an excitation wavelength at 1064 nm that is virtually free from the optical screening by the dark sheaths. The molecular imaging results give insights into not only secretion mechanisms of the sunscreen pigment but also interdependence between photosynthesis, nitrogen fixation, and redox homeostasis in one of the simplest forms of multicellular organisms.

## Introduction

Oxygenic photosynthesis by ancient cyanobacteria resulted in the rise of atmospheric oxygen concentration 2.3 – 2.4 billion years ago (Great Oxidation Event, GOE) ^1,2,3^. The oxygen gas is produced by the water oxidation in photosystem II (PSII), which has been preserved from the ancient cyanobacteria to modern chloroplasts in plants and algae. In spite of formation of an ozone layer after the GOE, UVA (320 – 400 nm) has continued to reach the surface of the earth^4^ and easily damaged PSII^5^. The repair mechanisms for PSII are deteriorated by reactive oxygen species (ROS) that are inevitable byproducts of photosynthetic electron transports in the presence of oxygen, especially under strong light^6^. The ancient cyanobacteria were thus threatened by the atmospheric oxygen, even though cyanobacterial respiration consumed some dioxygen. Some filamentous cyanobacteria secret a UV-screening pigment, scytonemin (Extended data Fig. 1), which is a yellow-brown secondary metabolite accumulating in the extracellular sheath^7,8^. The start of scytonemin biosynthesis has been estimated to be at least about 2.1 billion year ago^3, 9^. This timing and absorption peaks of scytonemin (~252, 278 and 386 nm)^8^ suggests that scytonemin became more indispensable after the GOE.

However, the biosynthesis of scytonemin are triggered by not only UV radiation, but also desiccation, and nutrient limitations^8^. Desiccation leads to an insufficient supply of electrons from water molecules and reduced molecular diffusions, which results in compromised photosynthetic electron transports generating ROS^10^. A strong radical scavenging activity of scytonemin has been reported,^11,12^ suggesting roles of scytonemin under various stress conditions generating radical species including some ROS. It remains unclear, however, whether the scytonemin at the extracellular sheath can scavenge the radicals generated by the photosynthetic electron transports in and around the thylakoid membranes in the intracellular regions^13^.

The oxidized scytonemin (OxScy) has been designated as “proper” scytonemin (Extended data Fig. 1)^7,14^. The other is reduced scytonemin (ReScy), which is the immediate precursor of the biosynthesis of OxScy^15^ and has been regarded to be limited to special reducing environments^7^. However, it has been pointed out that the ratio of ReScy/OxScy in an Antarctic lake sediment does not necessarily reflect the extracellular reducing conditions^16^. Differences between ReScy and OxScy have been shown in the growth suppression of human leukemia Jurkat cell^17^. The ratio of ReScy/OxScy, if measurable at a subcellular level, is expected to give a new insight into potential physiological roles of the redox transformation. Both OxScy and ReScy are virtually nonfluorescent^15^, while at least OxScy gives strong Raman signals^18,19,20,21^, but there has been no experimental Raman spectrum of ReScy. Here, we applied our newly developed Raman microscope with 1064 nm excitation (Extended data Fig. 2, Supplementary Fig. 1) to a diazotrophic filamentous cyanobacterium *Rivularia* M-261 ^22,23,24^, and visualized distributions of both OxScy and ReScy. Even when some visible light is shielded by the scytonemins, the high transmissions of both the excitation laser and Raman signals in the near infrared region enabled us to detect stoichiometric changes of intracellular photosynthetic pigments (mainly carotenoids bound to both PSII and photosystem I (PSI), and phycobilins bound to phycobilisomes), which reflect cellular differentiations. This study was partly aimed at understanding the polarity in the *Rivularia* filaments^22^, in which a single terminal cell at only the basal end becomes a nitrogen-fixing cell (heterocyst, Fig. 1a, Supplementary Fig. 2) and scytonemin localizations were also expected to be different from other scytonemin-producing cyanobacteria without obvious polarity^7, 21^. Here, we put emphasis on spectral differentiation of Raman signals of the two redox forms of scytonemin in cells and in their solid states, quantum mechanical predictions of scytonemin vibrations, and their subcellular distributions. The redox change of the scytonemin around the cells was found to be correlated with the heterocyst differentiations.

**Fig. 1.**
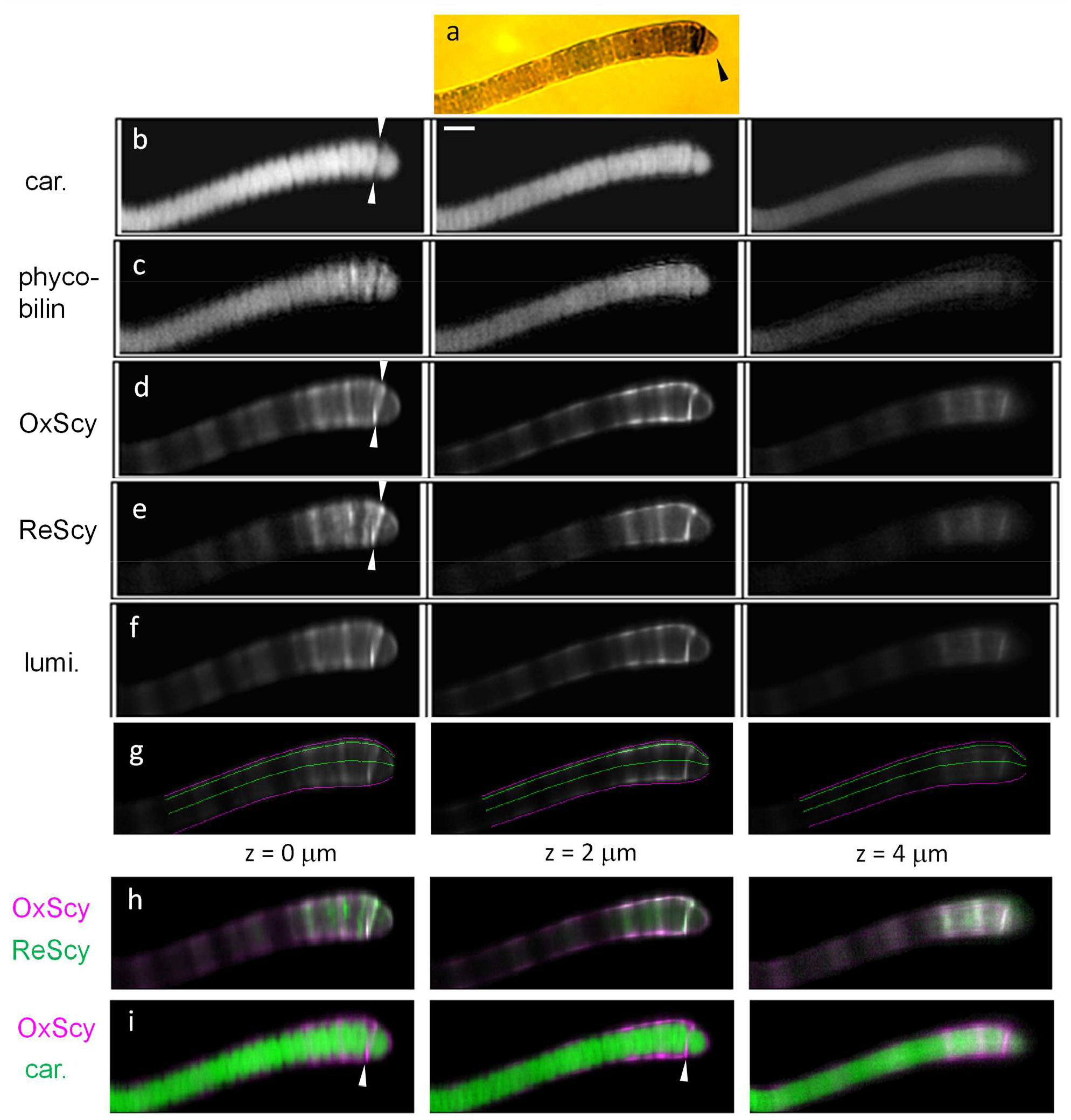
Images of a representative dark-sheathed *Rivularia* filament that contained the reduced scytonemin without the UVA treatment. The filament was classified as [w/ ReDS, no UVA]. (a) Bright field image. The black arrow head is pointing at a heterocyst at the terminal position. This end of the filament is called basal end and the other tapered end of the filament is called tip end (Supplementary Fig. 2). (b - e) Images of pigments derived from Raman scattering spectral data through singular value decomposition (SVD) and spectral fitting. Images of carotenoids (car.), phycobilins, oxidized scytonemin (OxScy) and reduced scytonemin (ReScy) are shown in (b), (c), (d), and (e), respectively. See Extended data Fig. 6 for the employed Raman spectra of the individual pigments. (f) Integrated intensities of autoluminescence overlapping with the Raman spectra in the detected wavelength range between 1266 and 1308 nm (corresponding to the Raman shift of 1500 – 1750 cm^−1^) are plotted. (g) Colored reference lines approximately parallel to the filament long axis (longitudinal axis) are superimposed on the gray scale image of OxScy in (d). The two magenta lines are drawn for outlining target cellular region for analysis, and the two green lines are drawn as examples of internally dividing points between the two magenta lines with internal division ratios of (53:47) and (91:9). (h, i) False-colored merged images for the pair of OxScy and ReScy (h) and the one of OxScy and carotenoids (i). The images of (b - i) were prepared for the three different focal planes at relative heights (z) of 0, 2 and 4 μm as shown between (g) and (h). The images at 0 μm show the lowest cross sections for the cells among the three relative heights. The white arrow heads in (b), (d), (e), and (i) indicate reference positions for the cell junction between the heterocyst and nearest vegetative cell. There are two points to be noted. First, ReScy is more internally located than OxScy at z = 0 and 2 μm in (h). See Supplementary Figs. 8 - 10 for filaments without substantial accumulatios of ReScy. Second, the heterocyst-vegetative cell junction is one of the hot spots for both ReScy and OxScy, as shown in (h) and (i).

## Results

### Definition of filament groups

On the UVA-treated preculture agar medium, both relatively short (young) and long (matured) filaments had dark sheaths extending from the basal end (heterocyst end) to nearly tip end (Supplementary Fig. 2). In contrast, when filaments were not treated with UVA, dark sheaths were often limited to the basal part of relatively long filament (including the terminal heterocyst and several vegetative cells). In this study, the regions of interest in the Raman microscopic imaging were selected to be only around the basal part (Fig. 1a). The Raman images for the same xy-range were obtained at three or five different depth positions (z) with an interval of 2 μm. The spectral analysis in the following sections led us to conclude that some dark-sheathed filaments (24 %) without the UVA treatment contained ReScy (Table 1, Extended data Table 1). Based solely on the spectral analysis and the preculture conditions before microscopic observations, we classified all analyzed filaments into four types (See also Methods): (i) non-UVA-treated filaments without dark sheaths ([w/o DS, no UVA]), (ii) UVA-treated filaments with dark sheaths but lacking ReScy-specific spectral features ([w/ OxDS, +UVA]), (iii) non-UVA-treated filaments with dark sheaths containing both ReScy and OxScy ([w/ ReDS, no UVA]), (iv) non-UVA-treated filaments with dark sheaths but lacking the ReScy-specific spectral features ([w/ OxDS, no UVA]). We first explain the spectral features specific to ReScy.

**Table 1.** Indexes of heterocyst differentiation and accumulation of the reduced scytonemin (ReScy) in the dark-sheathed filaments.

|  |  |  |  |
| --- | --- | --- | --- |
| Whether SVD spectra of the filaments contain the one characteristic to ReScy<sup>a</sup> (N: number of classified filaments) | Yes<br>(N=6)<sup>e</sup><br>[w/ ReDS,<br>no UVA] | No<br>(N=19)<sup>e</sup><br>[w/ OxDS,<br>no UVA] | No<br>(N=4)<sup>e</sup><br>[w/ OxDS,<br>+UVA] |
| Ratio of carotenoid-normalized phycobilin signals between heterocyst and connected vegetative cells.<sup>b,c</sup> Average (+/- SE) | 1.04*<br>(+/- 0.12) | 0.69<br>(+/- 0.06) | 0.232<br>(+/- 0.05) |
| Ratio of the integrated Raman signals between ReScy and carotenoid (ReScy/carotenoid) along the whole central longitudinal line of the filament.<sup>b,d</sup> Average (+/- SE) | 0.44*<br>(+/- 0.06) | 0.18<br>(+/- 0.03) | 0.17<br>(+/- 0.01) |

The asterisk (\*) indicates a statistically significant difference between [w/ ReDS, no UVA] and [w/ OxDS, no UVA] groups ( $p < 0.05$ , two-sided test).
(c): The ratio in this row was defined to be $\left( \frac{I_{\text{phycobilins}}}{I_{\text{carotenoids}}} \right)_{\text{het.}} / \left( \frac{I_{\text{phycobilins}}}{I_{\text{carotenoids}}} \right)_{\text{veg.}}$ ,
where, $I_{\text{phycobilins}}$ and $I_{\text{carotenoid}}$ indicate Raman signals of phycobilins and carotenoids at a pixel, respectively. The bracket with the subscript, ( )<sub>het.</sub> and ( )<sub>veg.</sub>, indicate that linear regions of averaging were selected to be inside the heterocyst and vegetative cells, respectively. The length of the linear regions was 1.6 – 5.2 $\mu\text{m}$ and 7.4 – 68 $\mu\text{m}$ for the heterocysts and vegetative cells, respectively.
(d): The ratio in this row was defined to be $(I_{\text{ReScy}})_{\text{whole}} / (I_{\text{carotenoids}})_{\text{whole}}$ ,
e) Distribution of individual data points are displayed in the correlation plot in Supplementary Fig. 3.

### Cellular Raman spectra showing OxScy and ReScy

Average single-cell spectra of the three groups, [w/ ReDS, no UVA], [w/ OxDS, +UVA], and [w/o DS, no UVA], showed qualitatively different Raman spectra (Fig. 2a, 2b, See Supplementary Fig. 5 for the selections of cell areas)^23^. In the dark-sheathed filaments, the cellular spectra showed scytonemin-specific Raman signals at around 1596, 1555, and 1172 cm^−1^ (Fig.2a),^19,25^ which were not detected in [w/o DS, no UVA] (Fig. 2b). Two Raman bands at around 1070 and 1365 cm^−1^ were prominent only in [w/ ReDS, no UVA] (Fig. 2a). The two bands were newly attributed to ReScy, which is further supported below. The Raman peak at 1555 cm^−1^ was substantially lower than that at 1596 cm^−1^ in [w/ OxDS, +UVA], while the two peaks showed comparable heights in [w/ ReDS, no UVA] (Fig. 2a). Although the distribution of scytonemin over the whole filaments and carotenoid-normalized average concentrations of OxScy in the basal regions in [w/ OxDS, +UVA] were more wide-spread and higher, respectively, than those in [w/ OxDS, no UVA] (Extended data Table 1), there was no clear difference in necessary component Raman spectra between the two groups.

**Fig. 2.**
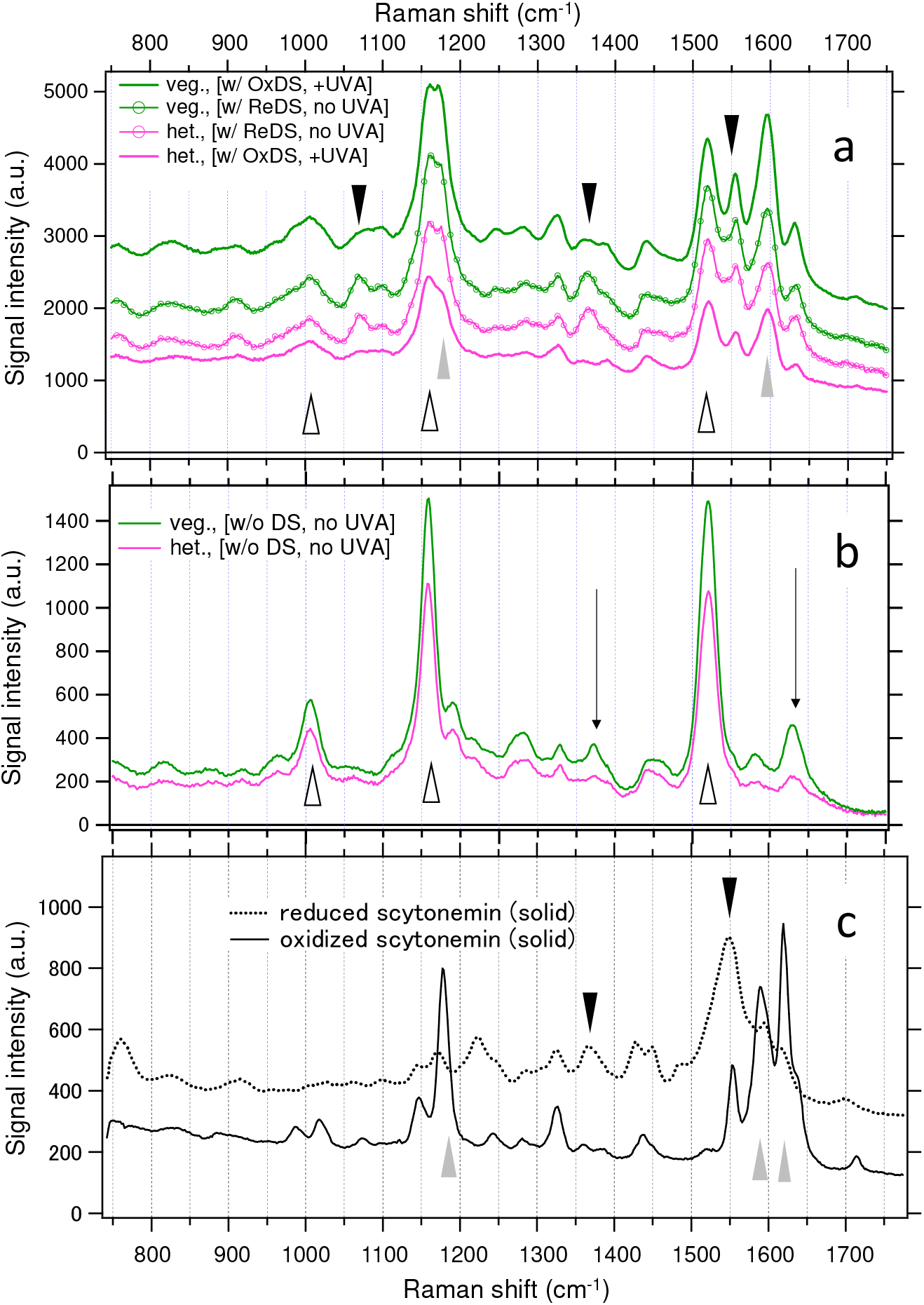
Representative Raman spectra to deduce spectral signatures of scytonemins. (a) Cell-type specific averaged single-cell spectra of *Rivularia* cells in the filament types of [w/ ReDS, no UVA] and [w/ OxDS, +UVA]. The plots in magenta and green represent spectra of heterocysts and vegetative cells, respectively in both (a) and (b). All spectra are averages of 3 cross-sections of different intracellular heights, and further averaged by 3 or 4 filaments of the equivalent cell positions and filament types. In addition, the spectra of the vegetative cells were averages of 5 connected vegetative cells nearest to the heterocysts. See Supplementary Fig. 5 for examples of the cell selections. The signal intensities reflect raw per-pixel intensities of the target single cells. (b) Cell-type specific averaged single-cell spectra of *Rivularia* cells in the filament type of [w/o DS, no UVA]. (c) Raman spectra of solid-state scytonemin under oxidized (solid line, OxScy) and reduced (dotted line, ReScy) conditions. In (a) – (c), the Raman bands at 1070, 1365 and 1555 cm are indicated by downward arrow heads (closed black), where ReScy in *Rivularia* cells shows characteristic Raman signals. Major scytonemin Raman bands that have been reported previously and attributable to OxScy are indicated by upward arrow heads (gray). Major Raman bands of carotenoids and phycobilins are indicated by upward open arrow heads and black straight lines with downward arrows, respectively.

### Raman spectra of solid scytonemin standard

The solid OxScy showed strong peaks at 1620, 1589 and 1177 cm^−1^ (Fig. 2c). The former two bands seem to correspond to the single Raman band at around 1596 cm^−1^ and the 3rd band seem to correspond to the one at 1172 cm^−1^ of OxScy in *Rivularia* (Fig. 2a). The three bands in the solid OxScy were significantly weakened in the solid ReScy, while two Raman bands at 1366 and 1550 cm^−1^ newly appeared in the solid ReScy and they also appeared in [w/ ReDS, no UVA] (Fig. 2a). These results support the view that ReScy accumulates in the filaments of [w/ ReDS, no UVA] and that accumulation of ReScy is far smaller than that of OxScy in the [w/ OxDS, +UVA] group. The ReScy-specific Raman band at 1365 cm^−1^ was sufficiently differentiated from the Raman band of phycobilins at around 1371 cm^−1^ (Fig. 2b).

### Spectral decompositions and pigment distribution maps

Singular value decompositions (SVD, Supplementary note 1) enabled us to know minimal spectral components to describe the Raman images of individual filaments (Supplementary Fig. 6, Extended data Figs. 3 - 5). Presence of ReScy was best manifested by the 5th SVD spectrum of a filament classified as [w/ ReDS, no UVA] (Extended data Fig. 3e), which showed concurrent negative ReScy-specific peaks at 1068, 1365, 1550 cm^−1^, as in the cellular spectra and/or in the solid ReScy (Fig. 2a, c). The obtained SVD spectra were fitted by weighted sums and/or differences of Raman spectra of photosynthetic pigments (carotenoids, phycobilins, and chlorophyll), the two redox forms of scytonemin (OxScy and ReScy) and luminescence (Extended data Figs. 3 - 6 and Supplementary note 2). Combinations of the fitting coefficients (e.g., Extended data Fig. 3-6), the singular values (e.g., Supplementary Fig. 6) and the scores of the spatial distribution of the major SVD components (e.g., Supplementary Fig. 7) yielded pigment distributions (e.g., Fig. 1, Extended data Fig. 7 for [w/ ReDS, no UVA], Supplementary Fig.8 for [w/ OxDS, +UVA], and Supplementary Figs. 9, 10 for [w/ OxDS, no UVA]). In the filament of [w/ ReDS, no UVA], carotenoids and phycobilins seemed to be rather homogeneously distributed in the intracellular regions over the whole filament (Fig. 1b, c), which reflect localizations of these pigments in the intracellular thylakoid membranes and their extensions over almost the whole individual cells (except centers of the cells when viewed with a higher spatial resolutions)^22^. In contrast, OxScy seemed to be concentrated at the extracellular sheaths (Fig. 1d, i, at 0 and 2 μm). Although ReScy seemed to be also concentrated at the extracellular sheaths to some extent (Fig. 1e, i, at 2 μm), there are more substantial concentrations of ReScy in the central regions of the filaments than OxScy (Fig. 1e, h, at 0 and 2 μm). The distribution map of the spectrally broad luminescence (Figs. 1f, 2a) was very similar to that of OxScy (Fig. 1d).

### Subcellular scytonemin distribution

In the plots of the pigment signals along the longitudinal line in the filament of [w/ ReDS, no UVA] (Fig. 1g, at z=0 μm, Fig. 3a, highest panel), there were high correlations among the OxScy, ReScy and luminescence. The particularly high correlation between OxScy and luminescence suggests that luminescence mainly comes from OxScy and/or some unknown molecule(s) colocalized with OxScy. If we concerned ourselves with pairs of peripheral and central regions with approximately the same pathlength from the basal end of the filament (Fig. 1g, h, z=0 μm, Fig. 3a, lowest panel), the local concentration ratio of ReScy/OxScy at a central point tended to be higher than that of the corresponding off-center point near the extracellular sheath.

**Fig. 3.**
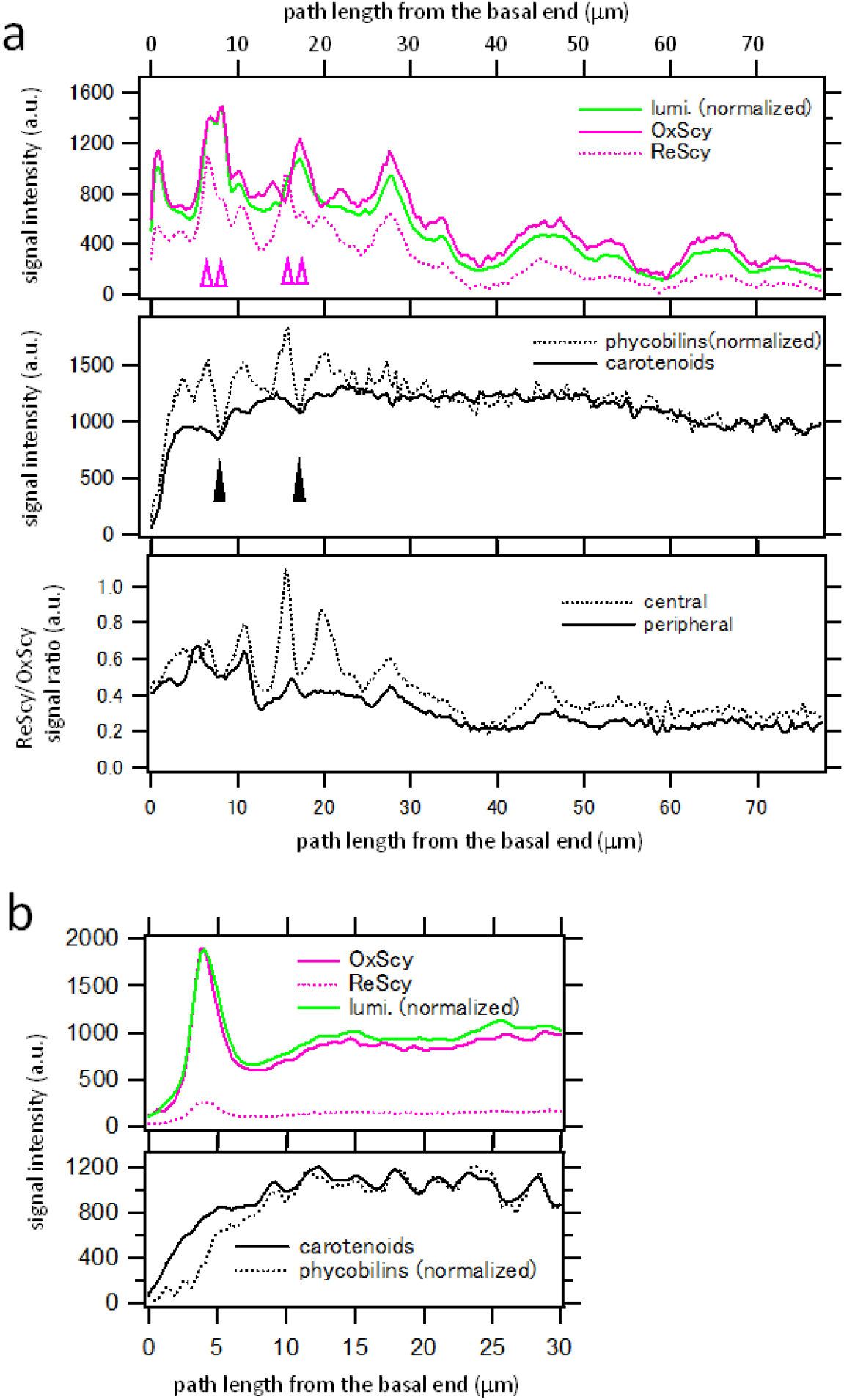
Pigment-Raman and autoluminescence signal profiles along filament long axes of the representative filaments. (**a**) The focal plane at z=0 μm of the filament type of [w/ ReDS, no UVA] shown in Fig.1 is analyzed here. Upper panel: profiles of scytonemins (ReScy and OxScy) and autoluminescence (lumi.) along the central longitudinal line (the green line with the internal division ratio of 53:47 between the two magenta lines, Fig. 1g). The intensity of the luminescence profile is normalized to be the same as that of OxScy at about 6.9 μm from the basal end. It should be noted that there may be multiphoton absorption of 1064 nm light by scytonemin and/or other pigments even if 1064 nm is not absorbed by one-photon absorption mechanism. The open magenta arrow heads show reference points at which the ratio in the Raman signals of OxScy and ReScy was varied significantly, showing uncorrelated behavior to some extent between the two components. Middle panel: profiles of carotenoids and phycobilins along the same central longitudinal line as in the upper panel. The intensity of the phycobilin profile is normalized to give the same average intensity as that of carotenoids between 30 and 50 μm from the basal end. The black arrow heads indicate positions of two cell junctions at which signals of carotenoids and phycobilins show local minima while those of scytonemins and luminescence show local maxima. Lowest panel: local ratios in the Raman signals between ReScy and OxScy that were estimated for relatively peripheral (around dark sheaths) and central regions at the image of z=0 μm of Fig.1. The peripheral region consists of two regions: one is between two lines with internal division ratios of (97:3) and (81:19), and the other is between two lines with internal division ratios of (22:78) and (6:94). The central region is between two lines with internal division ratios of (69:31) and (34:66). (**b**) The focal plane at z=2 μm of the filament type of [w/ OxDS, +UVA] shown in Supplementary Fig. 8 is analyzed here. The upper and lower panels of (b) are shown in analogous manners as in the upper and middle panels of (a), respectively. Although the inclusion of the ReScy in the spectral decomposition of the filaments treated with UVA ([w/ OxDS, +UVA]) was supported neither by the SVD spectra (Extended data Fig. 4) nor by the raw cellular spectra (Fig. 2a in the main text), the ReScy component was included here for a fair comparison of the filaments between the filament types of [w/ ReDS, no UVA] and [w/ OxDS, +UVA]. The resultant distribution of ReScy is very similar to that of OxScy (See merged image in Supplementary Fig.8j), indicating that accumulations of ReScy is probably negligible.

The heterocyst-vegetative (H-V) cell junction often showed a high concentration of scytonemins in all groups of [w/ ReDS, no UVA] (Fig. 1d,e,h,i, 3a, Extended data Fig. 7), [w/ OxDS, +UVA] (Fig. 3b, Supplementary Figs. 8, 12), and [w/ OxDS, no UVA] (Supplementary Figs. 9, 10). Some H-V and vegetative-vegetative (V-V) cell junctions were clearly observed as relatively dark transverse lines in the carotenoid and/or phycobilin Raman images (Fig. 1b, c, Extended data Figs. 7a,c, Supplementary Figs. 9, 10) or as local minima in the longitudinal profile of carotenoid and/or phycobilin signals (Fig. 3, Supplementary Fig. 11, 12).

Here we focus on the relatively well-resolved V-V cell junctions in one filament of [w/ ReDS, no UVA] (Extended data Fig. 7), where transverse cross sections at five focal heights show 3D distributions of the pigments as follows (Fig. 4). In the profiles overlapping with the cell junction (Fig.4a, Extended data Fig. 7d), the maximum of the ReScy appeared within 1.3 μm from that of carotenoid signal at three intermediate heights (z=−2, 0, +2 μm), while OxScy at the same heights showed two local maxima at more peripheral positions that correspond to sheath positions (further than 4.9 μm from the maximum position of carotenoids). Such qualitatively different distributions between ReScy and OxScy largely diminished if one examined the transverse profiles at higher (z=−4 μm) or lower heights (z=+4 μm). In the profiles overlapping with the center of a cytoplasmic space (Fig. 4b, Extended data Fig. 7b), the maximum position of ReScy at the intermediate heights (z=−2, 0, +2 μm) was at least 4.2 μm away from that of carotenoid maximum, while the signal ratios of ReScy/OxScy were lower than those near the cell junction (Fig. 4, z=0, −2 μm). Based on these results consistent with merged images (Extended data Fig. 7c), we propose that core regions and/or vicinities of core regions of some cell junctions of the filament type of [w/ ReDS, no UVA] seem to be hot spots of high concentrations of ReScy (Fig. 5).

**Fig. 4.**
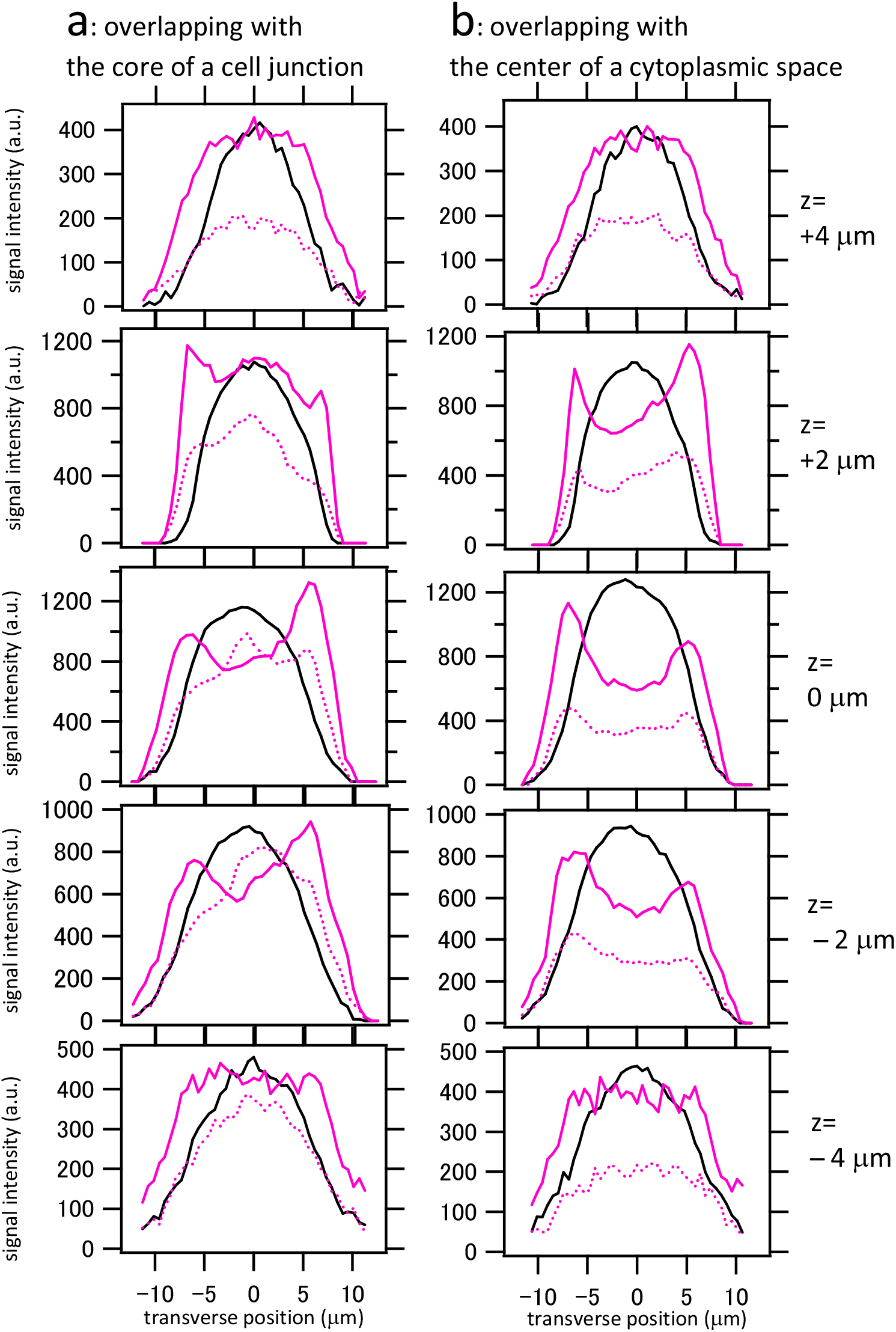
Raman signal profiles along transverse cross sections of a representative filament that contained the reduced scytonemin without the UVA treatment. This filament was classified as [w/ ReDS, no UVA]. The pigment profiles of OxScy (magenta solid line), ReScy (magenta dotted line) and carotenoids (black solid line) are plotted in the transverse cross sections indicated by the lines in Extended data Fig. 7d. (a) Plots along the transverse cross section overlapping with the core of a cell junction. (b) Plots along the transverse cross section overlapping with the center of a cytoplasmic space.

**Fig. 5.**
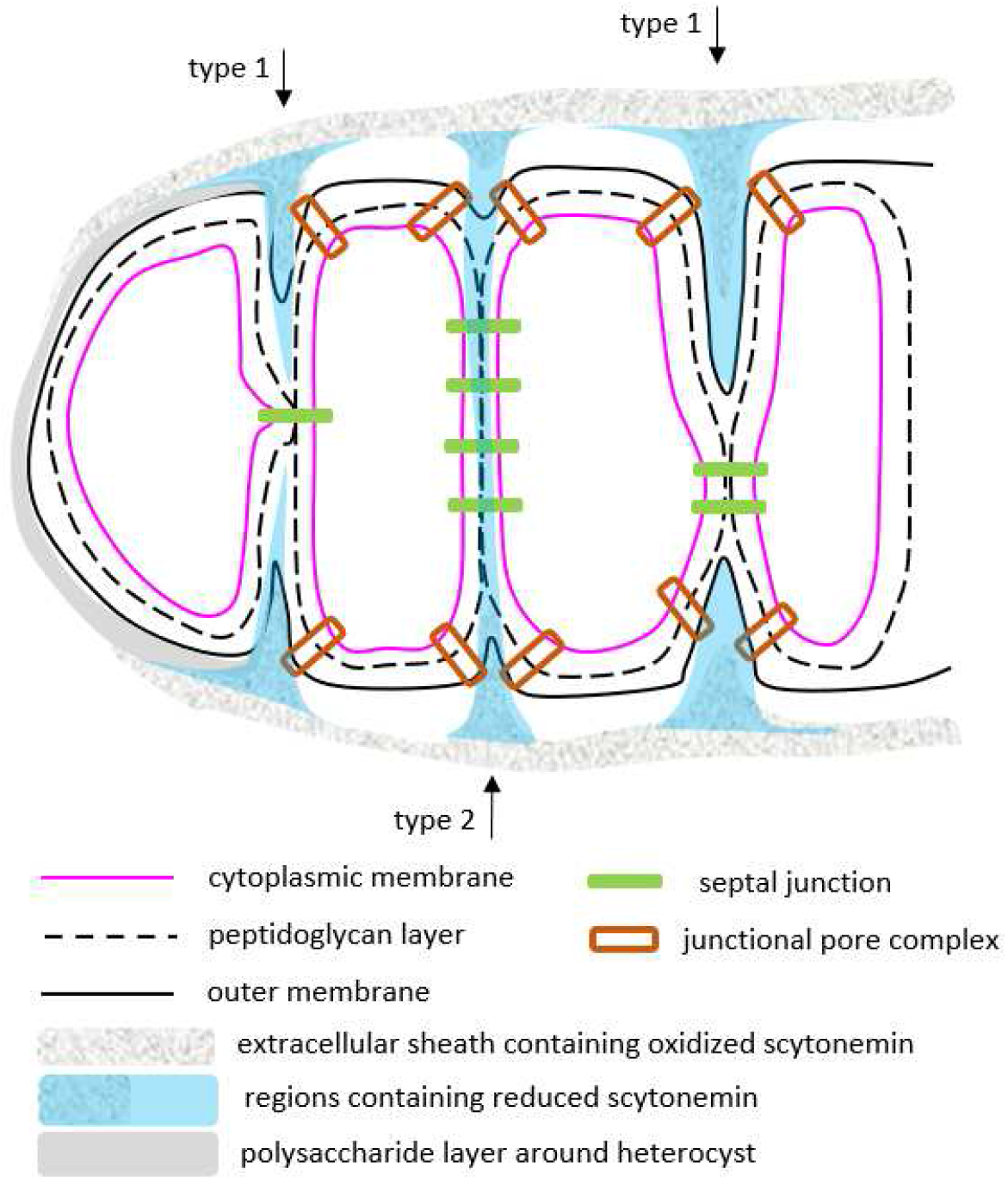
A model of the distributions of reduced and oxidized scytonemins (ReScy and OxScy) around the basal terminal of *Rivularia* M-261. The basic configurations of cytoplasmic membranes, peptidoglycan layers, outer membranes, polysaccharide layer around heterocyst, septal junctions, junctional pore complexes, and extracellular sheaths are largely following the models proposed in some previous works on other heterocyst-forming filamentous cyanobacteria^44, 45, 30, 46^. OxScy has been believed to be accumulated in the extracellular sheath, and it also showed substantial localizations near cell junctions in this work, which is in agreement with a previous report^21^. The distribution of ReScy that is significantly more internal than the one of OxScy may be realized by a deep recess of outer membranes near cell junctions, which prevents secreted ReScy from rapid oxidation (Type 1). Alternatively, some ReScy may be accumulated inside the periplasmic space near the cell junctions (Type 2). A third possibility is that some ReScy may reside in the cytoplasm, although our data alone are still unable to support or deny the last possibility. It should be noted that we have placed an empty space between “extracellular sheath containing oxidized scytonemin” and outer membrane of the cells, which may be possibly filled by scytonemin-free extracellular sheath. This feature is based on our observations that bright-field images show transparent regions between dark sheaths and cytoplasmic regions containing photosynthetic pigments in some dark-sheathed filaments (Fig. 1a, Extended Fig. 7b, Supplementary Figs. 2, 8 - 10).

The six filaments of [w/ ReDS, no UVA] (Table 1) showed clear accumulations of ReScy and/or OxScy near the core regions of the V-V cell junctions (e.g., Figs. 1,4, Extended data Fig. 7). It thus seems that ReScy remains unoxidized and accumulated near the core of V-V cell junctions because its transport to the extracellular sheaths were somehow stuck. In contrast, in all four filaments of the [w/ OxDS, +UVA] type and in the majority (>70 %) of nineteen filaments classified as [w/ OxDS, no UVA], we were able to find at least one z position (corresponding to a medium height for the cells) where central regions of V-V cell junctions did not show clear accumulations of OxScy (e.g., at z=0 and 2, in Supplementary Figs. 8, 9), in which scytonemins were predominantly localized at the extracellular sheaths and sufficiently oxidized. Some relatively minor filaments of the [w/ OxDS, no UVA] type (<30 %) showed substantial accumulations of OxScy near the core regions of V-V cell junctions (Supplementary Fig. 10).

### Quantum chemical predictions of scytonemin vibrations

The monomer moiety of scytonemin (MmScy, Extended data Fig. 1) is an intermediate synthesized in the cytoplasm, and it is transported to the periplasm (Fig. 5), where oxidative dimerization of MmScy yields ReScy that undergoes a rapid autooxidation to OxScy^15^. Given that ReScy was more internally located than OxScy in some *Rivularia* filaments, the question arises whether the Raman signals of ReScy that we postulated is attributable to MmScy or ReScy. We carried out quantum chemical calculations to predict Raman spectra of the OxScy, ReScy and MmScy (Extended data Fig. 8). These predicted that the intense Raman band of the OxScy around 1610 cm^−1^ shifts to 1559 cm^−1^ in the ReScy. This band is attributable mainly to the stretching vibration of the bond between two carbon atoms (C1 and C22) connecting the two molecular halves of OxScy and ReScy (Extended data Fig. 1, Supplementary Fig. 13). It is noteworthy that the angles of displacement vectors of C1 and C22 atoms relative to the C1-C22 bond is significantly altered from OxScy to ReScy (Supplementary Fig. 13), which is caused by the mixing with the bending motion of the N-H bonds that only exist in the ReScy and it results in the low-frequency shift of 51 cm^−1^. The theoretical band positions at 1610 cm^−1^ (OxScy) and 1559 cm^−1^ (ReScy) seem to be in reasonable agreement with the experimental ones at 1596 cm^−1^ (OxScy) and 1555 cm^−1^ (ReScy) in the *Rivularia*, respectively (Fig. 2a, Extended data Fig. 6). On the other hand, only three strong Raman bands at 1593, 1598 and 1616 cm^−1^ were predicted for MmScy between 1500 and 1700 cm^−1^ (Extended data Fig. 8, Supplementary Fig. 14), for which there are three corresponding vibrational modes of OxScy at nearly the same frequencies (Extended data Fig. 8). Thus, the new scytonemin-associated Raman signals that were found to be more internally localized than OxScy are attributable not to MmScy, but to ReScy. We found five vibrational modes that are potentially attributable to the ReScy-specific Raman band at 1365 cm^−1^ (Figs. 2a, c, Supplementary Fig. 15.)

### Cellular differentiation and scytonemin redox conditions

The matured heterocysts in the filaments without dark sheaths of *Rivularia* M-261 (as [w/o DS, no UVA]) have been reported to show lower phycobilin/carotenoid ratios than the nearby vegetative cells (Fig. 2b)^23^. However, the filaments of the [w/ ReDS, no UVA] type tended not to show a decrease of phycobilins/carotenoids ratio in their heterocysts (Table 1, Fig. 2a, 3a, Supplementary Figs. 3, 11, 16), which possibly means that they are not (yet) heterocysts but just basal terminal cells. In the same agar medium in which we found the filaments of [w/ ReDS, no UVA], most dark-sheathed filaments were grouped as [w/ OxDS, no UVA], in which heterocysts tended to show lower phycobilin/carotenoid ratios than the nearby vegetative cells (Table 1, Supplementary Fig. 3). In addition, the four filaments of the [w/ OxDS, +UVA] type also showed significant decreases of phycobilins/carotenoids ratio in their heterocysts in comparison with the vegetative cells (Fig. 3b, Supplementary Figs. 3, 12 and 16).

## Discussion

Not only extracellular sheaths^12,26^ but also cell junctions of filamentous cyanobacteria seem to be occupied by scytonemins, as found by a previous study applying coherent anti-Stokes Raman scattering (CARS) microscopy to UVA-treated *Nostoc commune* that probably contained only OxScy^21^. We confirmed a similar tendency in non-UVA-treated *Rivularia* M-261 by our spontaneous Raman scattering images, in which signals are more faithfully proportional to microscopic molecular concentrations than CARS^27^ (Figs. 1, 4, Extended data Fig. 7). Our 3D Raman images have further shown that ReScy tended to be accumulated near the core regions of the cell junctions (Figs. 4, 5), which are more internal than those of OxScy.

Some ReScy may accumulate in extracellular sheaths invaginating in deep recesses (constricted regions) of outer membranes at cell junctions (Type 1, Fig. 5). Although this effect alone may explain the ReScy/OxScy accumulations at the H-V cell junctions where constrictions of the cell widths are typically more conspicuous than V-V cell junctions (Figs. 1, Extended data Fig. 7, Supplemental Figs. 8 - 10), it seems insufficient to make the peak of ReScy so close to the core regions of the V-V cell junctions. Accumulations of ReScy inside the periplasmic space near the cell junctions (Type 2, Fig. 5) seems more convincing, because the oxidative dimerization from the MmScy to ReScy takes place in the periplasm^15^. In any scenario, our results suggest that sites of secretion of MmScy and/or ReScy are near the core of the cell junction and that some ReScy can remain unoxidized around those sites. The accumulated ReScy in the periplasmic space may work as an antioxidant for both the periplasmic space and cytoplasm, which may help *Rivularia* survive under stressful conditions. Secretion sites of scytonemin have not been elucidated^28^, while those of extracellular polysaccharides, major components of extracellular sheaths^29^, have been reported to be junctional pore complexes surrounding the septal junctions at least in the motile phase of scytonemin-producing *Nostoc punctiforme* (Fig. 5)^30^. Secretions of scytonemins may be coordinated with that of extracellular polysaccharides partly through closeness between their secretion sites.

Substantial decrease in the ratio of phycobilin/carotenoid compared to that of vegetative cells is a typical feature of matured heterocysts including those of *Rivularia* M-261^22, 23^. This reflects the decrease of light-harvesting phycobilisomes (binding phycobilins) and PSII (binding carotenoids) for the anaerobic conditions required for the heterocysts, which results in no fixation of carbons in mature heterocysts^31^. On the other hand, the amount of PSI (binding carotenoids) of the heterocyst remains as high as those in vegetative cells^22^, which carries out a cyclic electron transport to produce ATP for nitrogen fixation^31^. The filaments containing ReScy often lacked the heterocyst-specific spectral features at their basal terminal cells (Fig.3, Table 1, Supplementary Figs. 3, 16). The perturbation of heterocyst formation by the abnormal redox conditions seems likely, considering that some cross-talks between oxidation stress responses and heterocyst formations have been reported^32,33, 34^. On the other hand, this may possibly be a sign of another cellular differentiation (e.g., dormant cells (akinetes)) toward survival of the cells under desiccation and/or starvation^34^. It is to be noted that *Rivularia* M-261 is probably equipped with some desiccation-tolerant mechanisms, because it was originally found from a cement wall in Bangkok, Thailand^24^, and we often succeeded to recover phototrophic growth of filaments inoculated from agar cultures that were more desiccated than those analyzed in this study. The physiological effects of ReScy need to be further clarified by a long-term time-lapse observation of desiccation and rewetting of individual filaments after Raman imaging of OxScy and/or ReScy. It is also noteworthy that the basal end of basal terminal cell (heterocyst) in some dark-sheathed filaments is not substantially surrounded by the dark sheaths, while H-V cell junction is often a hot spot of ReScy and/or OxScy (Fig. 3b, Extended data Fig. 7, Supplementary Fig. 2, 8 - 10, 12). This probably reflects the absence of secretion sites of scytonemin on the basal terminal cell (heterocyst) and no significant transport and/or diffusion of scytonemins toward the basal terminal end, which is reasonable given that UVA-sensitive PSII is absent and there is no ability to fix carbons in mature heterocysts.

In conclusion, at least relatively long durations of preculturing caused accumulations of ReScy near core regions of cell junctions in some *Rivularia* filaments probably through desiccation and/or starvation even when UVA was not illuminated, while the UVA treatment alone on a relatively fresh preculture did not immediately lead to a substantial accumulation of ReScy. The accumulation of ReScy was correlated with the differentiation status of the basal terminal cell, which implies importance of redox homeostasis of the cells for their differentiations. The spontaneous Raman scattering imaging with a 1064 nm laser excitation and simultaneous broad spectral coverage is thus useful for finding hitherto unreported Raman spectra of metabolites and their subcellular distributions without staining, by which we will hopefully understand various physiological states and survival strategies of multicellular prokaryotes even in relatively deep regions where visible light cannot penetrate due to thickness and/or shielding of visible light (e.g., Supplementary Fig. 17).

## Methods

### Preparations of cyanobacteria! cells

*Rivularia sp*. IAM M-261, which is phylogenetically very similar to *Calothrix sp*. PCC 7715^35^, was obtained from the culture collection of Institute of Molecular and Cellular Biosciences, the University of Tokyo (IAM Collection). The strain has been transferred to Microbial Culture Collection at the National Institute for Environmental Studies (NIES Collection Tsukuba, Ibaraki, Japan)^22, 24^. *Rivularia* filaments were grown on agar media based on the exogenous-nitrogen-free BG-11 (BG110) prepared by removing NaNO3 and replacing ferric ammonium citrate with ferric citrate in a glass tube (Supplementary Fig. 17). A white fluorescent lamp was used without dark periods at a flux density of photosynthetic photons of about 10 μmol photons m^−2^ s^−1^ at the glass tube position in an incubator at 28 °C. Before harvesting the cells for microscopic observations, the *Rivularia* cells were precultured in the BG11_0_ media in a polystyrene petri dish in the same incubator for 20 - 35 days (Supplementary Fig. 18).

For the UVA-treated *Rivularia* filaments with dark sheaths containing only OxScy and almost no ReScy ([w/ OxDS, +UVA]), UVA from two LED light bulbs with center wavelengths of 369 and 371 nm was additionally illuminated with a total power of about 1 mW/cm^2^ at the sample position for the last three days of the preculture period (27 days), and four filaments with dark sheaths were arbitrarily selected for Raman microscopy. For the other filament types, no UVA illumination was added. We did not find dark sheaths in most of the filaments after the preculture period of 20 days, and three filaments without dark sheaths were arbitrarily selected for Raman microscopy ([w/o DS, no UVA]). Even when there was no UVA, filaments having dark sheaths were often found after 31 – 35 days of the preculture period. The relatively long preculture period resulted in a shrinkage of the agar layer in the preculture dish, suggesting desiccation and/or nutritional starvation stress(es) to the *Rivularia* cells (Supplementary Fig. 18). From the surface of such agar media, we arbitrarily selected 25 filaments having dark sheaths for Raman microscopy (Extended data Table 1). Based on whether spectral signatures of ReScy was found or not (See Result sections), six and nineteen filaments were classified as [w/ ReDS, no UVA] and [w/ OxDS, no UVA], respectively (Table 1). We avoided observing short motile filaments (hormogonia). All selected filaments had a full length of at least 88 μm or longer, and they also showed a clear polarity in that several vegetative cells near the basal end (heterocyst end) were wider than those of the other end that is often described as tapered end (Supplementary Fig. 2).

### Micro-spectroscopic Raman imaging

The basic setup was the same as previously described^23^, but the excitation laser wavelength was changed into 1064 nm and a linearly elongated illumination with a length of about 66 μm was newly employed (Extended data Fig. 2). The average total laser power at the sample position was 340 mW. The whole region illuminated by the 1064 nm laser can generate Raman scattering signals that were detected by an InGaAs camera (640 × 512) attached to an imaging polychromator. The images of Raman signals consisted of signals from different spatial positions (about 200 pixels in the vertically arranged 512 pixels) and different vibrational frequencies (different wavelengths, along horizontally arranged 640 pixels). After corrections of pixel defects, cosmic ray noise and image distortions including wavelength calibrations, one single camera image yielded Raman spectra at 190 positions with an interval of 0.33 μm (Y axis, 190×0.33 μm ~63 μm). An XYZ-piezo stage holding the glass-bottomed dish with cyanobacterial cells was moved along axes perpendicular to the 1064 nm laser line with step sizes of 0.33 μm (X axis) or 2.0 μm (Z axis), and the stepping motion was in synchrony with the camera exposure. The full width at half maximum of the point-spread function (spatial resolution) of the setup was estimated to be 0.51, 1.1, and 6.0 μm for the X, Y and Z axes, respectively (Supplementary Fig. 1).

For the Raman microscopic imaging of *Rivularia* cells, a piece of agar block was transferred from the preculture agar plates to a cover-glass-bottomed dish for observing cells on the agar surface (Supplementary Fig. 18). Formation of air gaps between the cover glass and the cells was avoided by placing a drop of distilled water before the transfer of the agar piece. Accumulation time at a single XZ position (single line along Y axis) was 10.0 s for the imaging of *Rivularia* cells, by which effective time to acquire single pixel Raman spectrum was about 0.053 s (~10 s/190).

### Image and spectral analysis

Singular value decompositions (SVDs) and the subsequent spectral decompositions were carried out by our in-house programs based on Mathematica (Wolfram research). SVD was applied to individual *Rivularia* filaments separately. In each data set containing Raman spectra at many different 3D points of a single filament, only pixels with substantial Raman/luminescence signals were selected based on a threshold and they were used as the target region for the SVD analysis. The details of the procedures are described in Supplementary Notes 1 and 2. The bright field images were auto-contrasted in XnView software.

### Raman spectra of oxidized and reduced forms of scytonemin standard

Scytonemin from naturally growing *Nostoc commune* colonies was used as a standard^11^. The solvent (DMSO) was removed by a low-pressure evaporation, and the solid-state scytonemin was placed on the cover-glass in a glass-bottomed dish, which first gave Raman spectra of the oxidized scytonemin (OxScy). An average Raman spectrum of OxScy (Fig. 2c) was calculated from those measured at nine different microscopic regions of the solid-state scytonemin. After the measurements of OxScy, an aqueous solution of sodium dithionite (5 mM) was added to the glass bottomed-dish in order to reduce scytonemin chemically, and the dish was tightly sealed with a cap by a sealing film (Parafilm) of a few layers. An average Raman spectrum of the reduced scytonemin (ReScy) was estimated from those in the same nine regions (Fig. 2c). It took about 3 hours during the Raman spectral imaging of the nine regions, in which average Raman spectrum of an individual region was not distinguishable from the others.

### Theoretical calculation of Raman spectra of scytonemins

Normal mode analysis was performed on ReScy, OxScy, and MmScy molecules isolated in the gas phase using the DSD-PBEP86 double-hybrid DFT^36^ with the def2-SVP basis sets^37^. The Raman activity^38^ for each normal mode was evaluated by numerical differentiations of the analytic dipole polarizability with the domain-based local pair natural orbital approximation ^39, 40^ at the equilibrium geometry. All the calculation were performed with the ORCA5 software package ^41^. We selected a vibrational frequency scaling factor of 0.943 that seemed to best reproduce the experimental Raman spectra, and the selected value was close to the values (0.9548 or 0.9605) recommended by theoretical groups using very similar calculation methods to ours^42,43^.

## Data availability

Source data are provided with this paper. All other data are available from the corresponding author upon reasonable request.

## Code availability

The scripts used to perform curve fittings of SVD spectra by sum of pigment Raman spectra are provided with this paper. The software package ORCA 5.0 used to perform the normal mode analysis is available from the ORCA web site (https://orcaforum.kofo.mpg.de). All other codes are available from the corresponding author upon reasonable request.

## Acknowledgements

We thank Profs. M. Terazima (Kyoto Univ.), T. Shiina (Setsunan Univ.), Y. Kimura (Doshisha Univ.), H. Oh-Oka (Osaka Univ.) and M. Katayama (Nihon Univ.) for helpful scientific advice. This work was supported in part by the Japan Science and Technology Agency (the Precursory Research for Embryonic Science and Technology, “Chemical conversion of light energy”, to S.K.), MEXT-Supported Program for the Strategic Research Foundation at Private Universities (2015 - 2018, no. S1511025 partly to S.K.), “Molecular Science for Supra-functional Systems” (Area No.477, 19056012 to S.K.), JSPS KAKENHI (17H03968 partly to S.K. and 18K06153 partly to S.K.), the Murata Science Foundation (to S.K.), the Kyoto University Foundation (to S.K., in the fiscal year of 2021) and Imaging Science Project of the Center for Novel Science Initiatives (CNSI), National Institutes of Natural Sciences (NINS) (no.IS281001 to S.K.).

## Author contributions

S. N., K.T. and S.K. found growth conditions of dark-sheathed *Rivularia* without UVA treatments; S.K. designed the line-scanning Raman scattering spectral microscopy; K.T. and S.K. constructed the Raman microscope and designed experiments; T. S. extracted the oxidized scytonemins from *N. commune*, K.T. performed the Raman spectral imaging of *Rivularia* and solid-state scytonemins; Y.K. performed the calculations of Raman spectra of the scytonemins; K.T. and S.K. analyzed the results; K.T. and S.K. wrote the manuscript with comments from all authors. All authors read and approved the final manuscript.

## Ethics declarations

### Competing interests

The authors declare no competing interests.

---

**Extended data Fig. 1.**
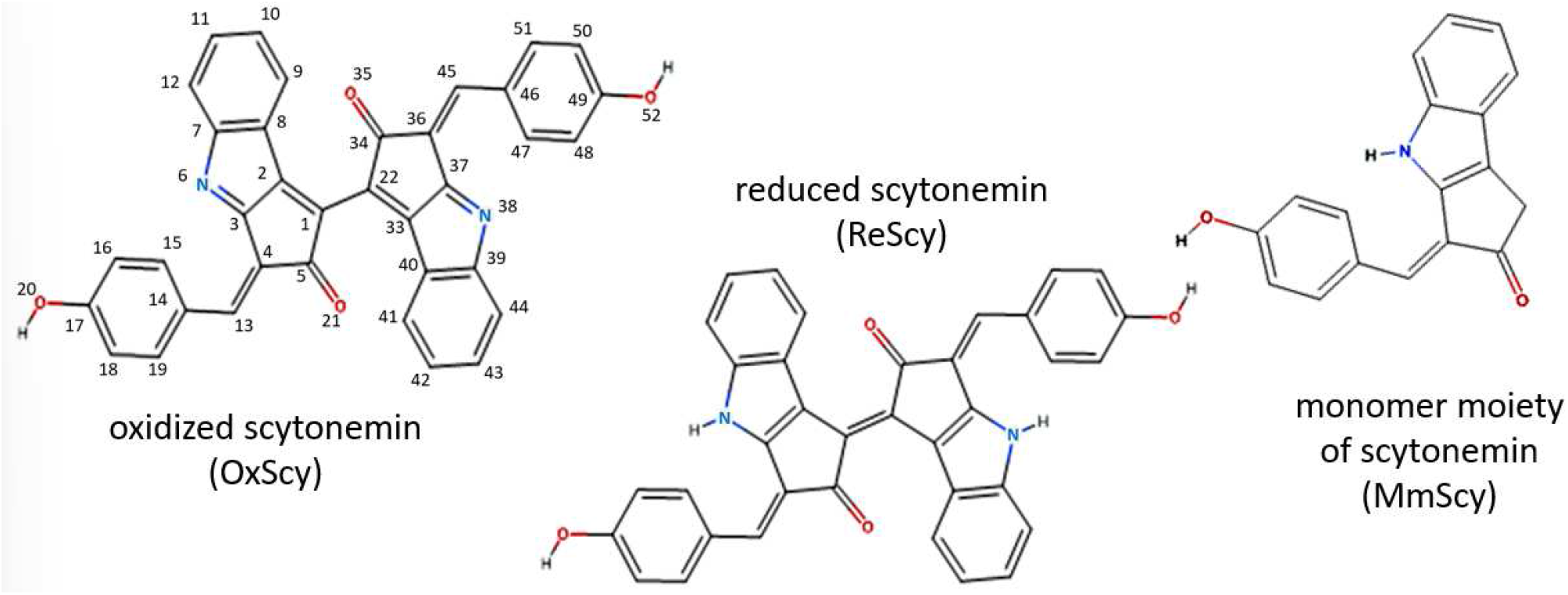
Molecular structures of the scytonemins (OxScy and ReScy) and their biological precursor form (MmScy). Atoms in OxScy are numbered according to a reference^25^. Only carbon, oxygen and nitrogen atoms are labeled here, and the same numbering system is applicable to ReScy.

**Extended data Fig. 2.**
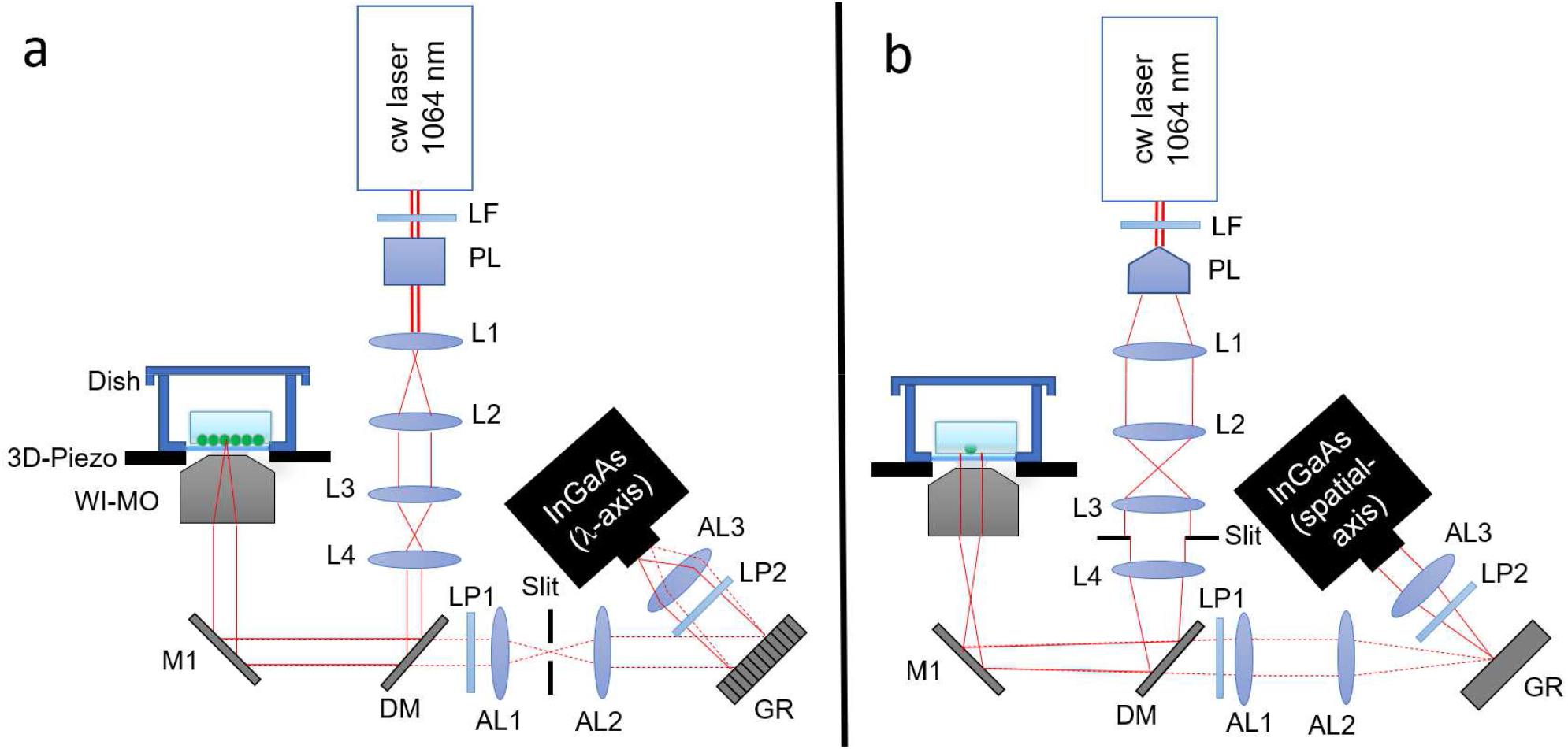
Schematic diagram of the home-made line-scanning Raman spectral microscope. a) Viewing angle to see the spectral detection of a focus point in a sample. b) Viewing angle to see the linear image detection of a focus line in a sample. The basic setup was the same as previously reported^23^. WI-MO: water-immersion microscope objective lens, AL1-AL3: achromatic lenses for the near infrared light, L1-L4: lenses (L1: f=15 mm, L2: f=100 mm, L3: f=200 mm, L4: f=300 mm), DM: long-pass dichroic mirror with an edge wavelength of 1080 nm, LP1 and LP2: long-pass filters with an edge wavelength of 1085 nm, M1: flat mirror, 3D-Piezo: piezo actuator stage to move the sample positions in the three dimensions, Dish: glass-bottomed dish, GR: diffraction grating, PL: Powell lens (fan angle = 10 degrees), LF: laser line filter (at 1064 nm).

**Extended data Table 1.** Basal-region-area-averaged pigment Raman signals and their ratios in the three groups of dark-sheathed *Rivularia* filaments.

| [filament type]<br>N=number of filaments | [w/ OxDS, +UVA]<br>N=4<sup>d</sup> | [w/ OxDS, no UVA]<br>N=19<sup>d</sup> | [w/ ReDS, no UVA]<br>N=6<sup>d</sup> |
| --- | --- | --- | --- |
| carotenoids<sup>a</sup>(a.u.) | 557 (±32) | 449 (±23) | 502 (±35) |
| OxScy<sup>a</sup> (a.u.) | 1310* (±169) | 530(±54) | 609 (±54) |
| OxScy/carotenoids<sup>b</sup> (a.u.) | 2.34*(±0.27) | 1.26(±0.15) | 1.27(±0.19) |
| ReScy<sup>a,c</sup> (a.u.) | 250 (±50) | 168 (±18) | 321* (±46) |
| ReScy/carotenoids<sup>b,c</sup> (a.u.) | 0.44(±0.08) | 0.40(±0.05) | 0.64*(±0.08) |
| ReScy/OxScy<sup>b,c</sup> (a.u.) | 0.19(±0.02) | 0.32(±0.01) | 0.55*(±0.10) |

The asterisk (\*) indicates a statistically significant difference between the marked group and combination of the other two groups ( $p < 0.05$ , two-sided test).
(a) These values were average signals in the basal regions of filaments given as follows. In each filament, maximum signals of four pigments (carotenoids, phycobilins, OxScy and ReScy) were separately determined (See Supplementary Note 2). If at least one pigment Raman signal at a pixel exceeded 10 % of its maximum signal in the same filament (threshold), the pixel was included in the basal region. The basal-region-area-averaged pigment Raman signal of a filament was thus given by the average signal in the given basal region. Thus, the averaged area included both intracellular regions and extracellular sheaths.
(b) These values were averaged ones over multiple filaments, for which ratios between the basal-region-area-averaged pigment Raman signal of individual filaments were obtained.

**Extended data Fig. 3.**
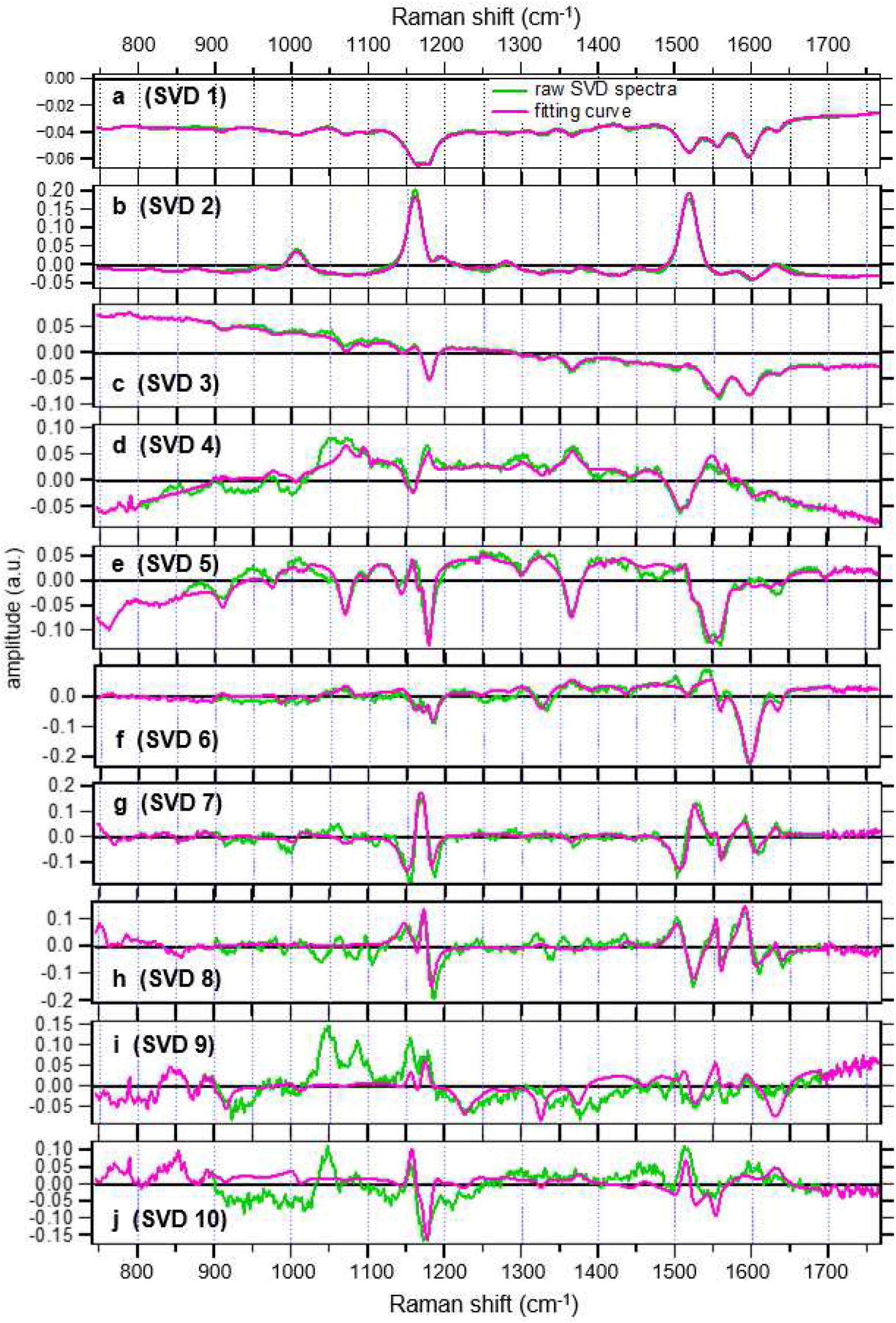
Major SVD spectra for the representative filament that contained the reduced scytonemin without the UVA treatment. This filament was classified as [w/ ReDS, no UVA] type. The SVD spectra in green are placed in the descending order of the singular values plotted in Supplementary Fig. 6a. Magenta plots indicate best fits given by the sum and/or differences of autoluminescence spectra and the 11 pigment Raman spectra shown in Extended data Fig. 6 (See also Supplementary note 2). The target filament is identical to that shown in Fig 1 in the main text.

**Extended data Fig. 4.**
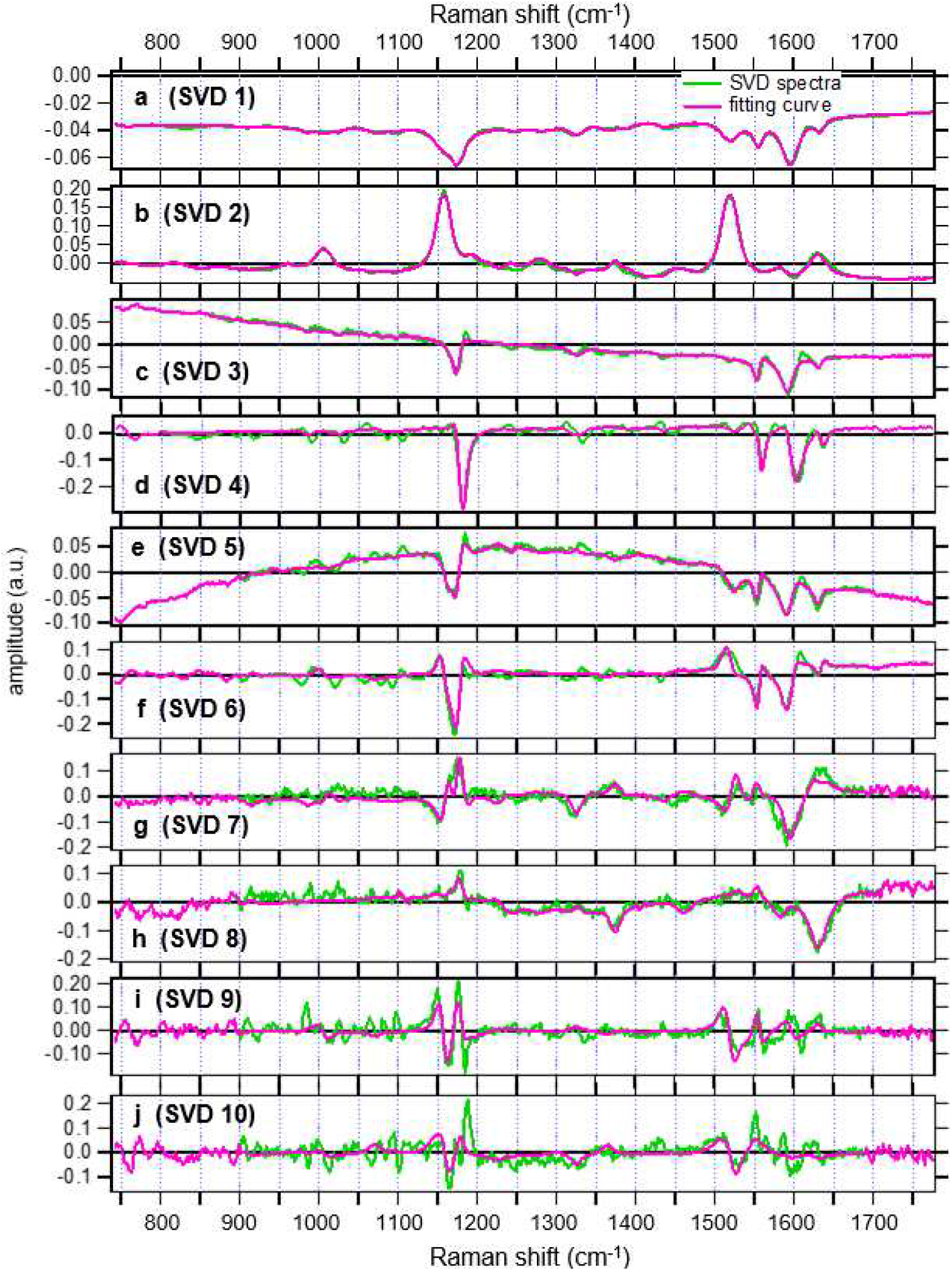
Major SVD spectra for the representative filament that had dark sheaths without the reduced scytonemin after the UVA treatment. This filament was classified as [w/ OxDS,+UVA] type. The SVD spectra in green are placed in the descending order of the singular values in Supplementary Fig. 6b. Magenta plots indicate best fits given by the sum and/or differences of autoluminescence spectra and the 11 pigment Raman spectra shown in Extended data Fig. 6 (See also Supplementary note 2). The target filament is identical to that shown in Supplementary Fig 8.

**Extended data Fig. 5.**
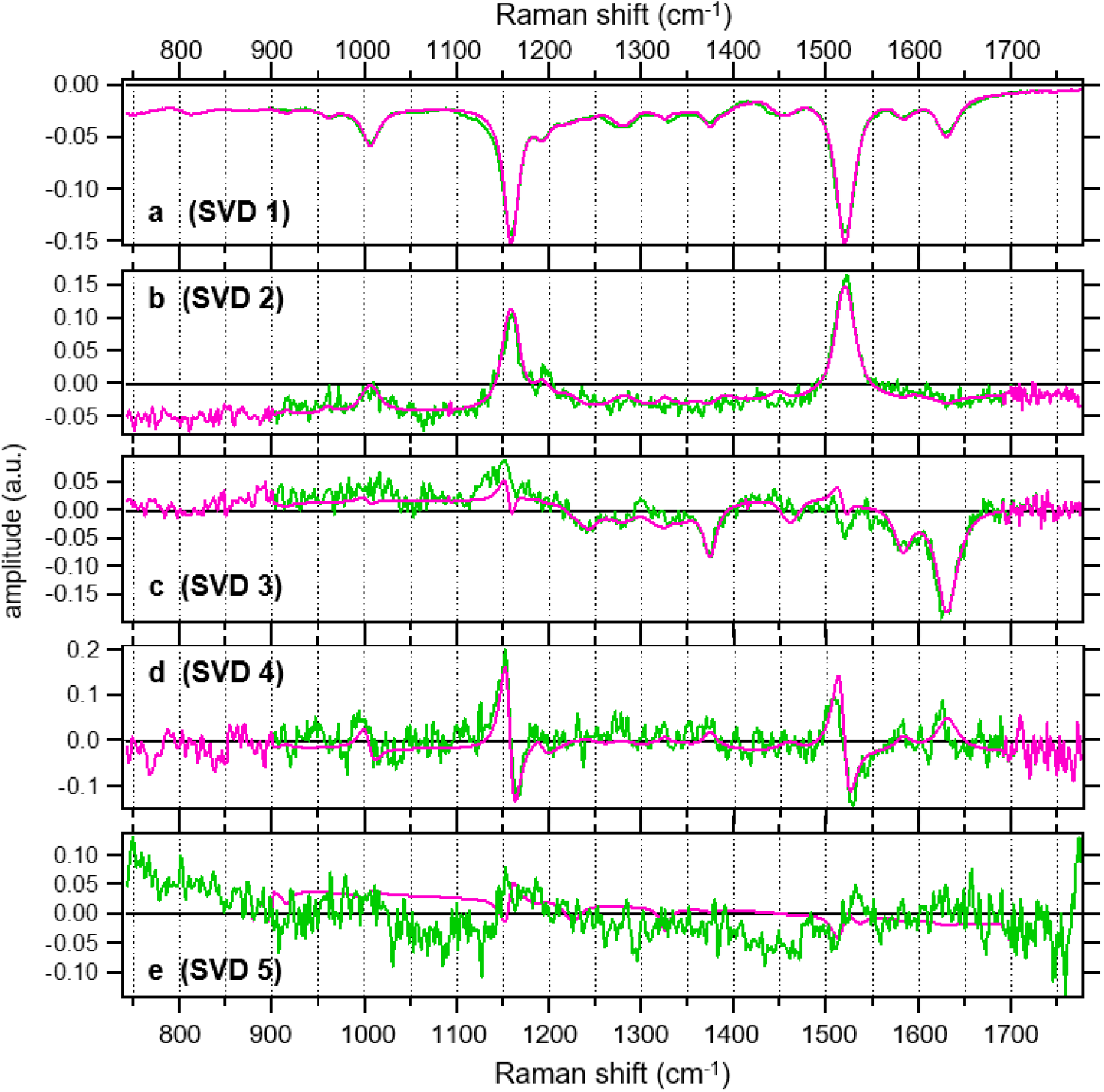
Major SVD spectra for the representative filament that had no dark sheath. This filament was classified as [w/o DS, no UVA]. The SVD spectra in green are placed in the descending order of the singular values in Supplementary Fig. 6c. Magenta plots indicate best fits given by the sum and/or differences of the 11 pigment spectra shown in Extended data Fig. 6 and autoluminescence spectra (See also Supplementary note 2).

**Extended data Fig. 6.**
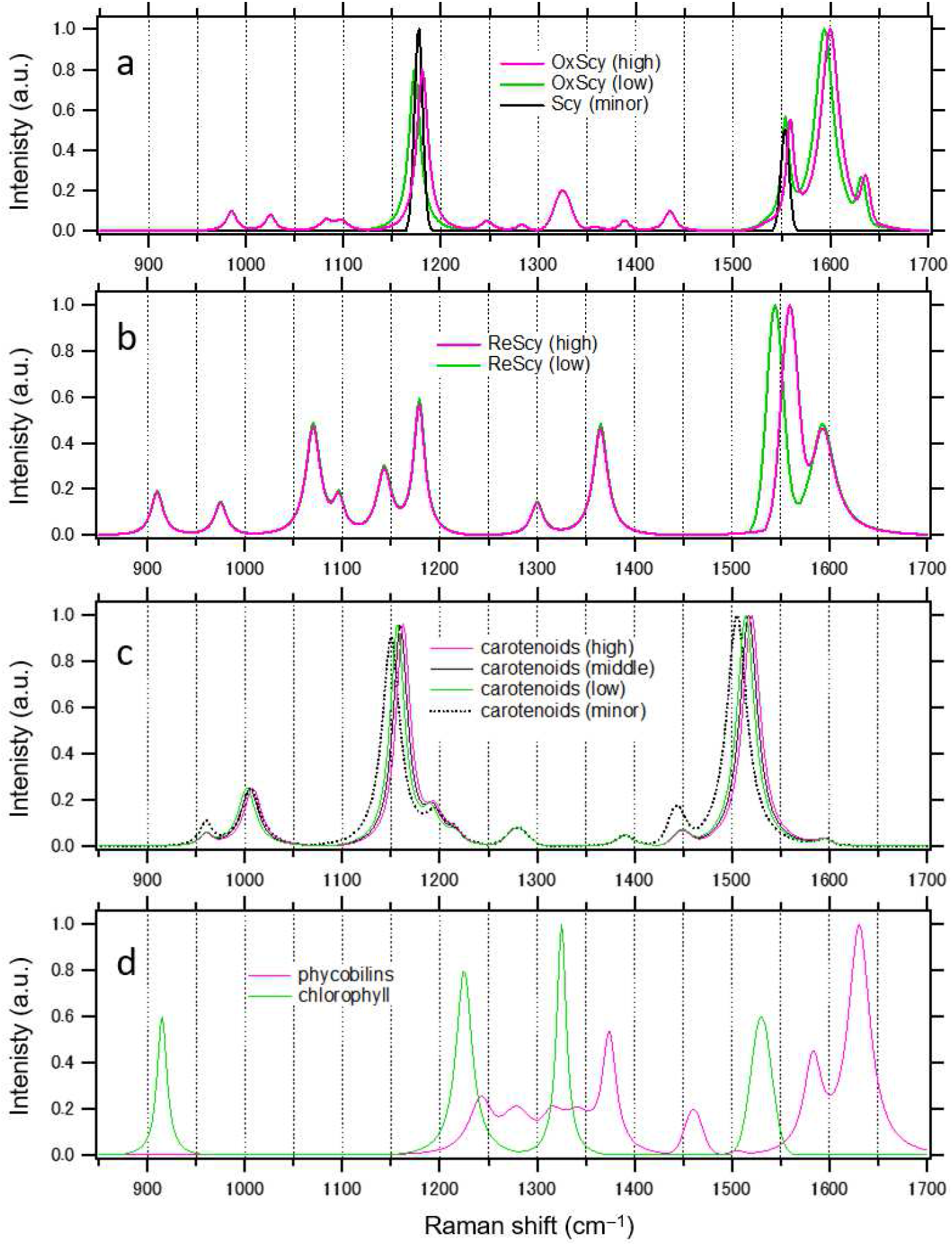
Raman spectra of the pigments used for the SVD spectral fitting. The fitting results are shown in Extended data Figs. 3 - 5. Multiple spectral forms were assumed for oxidized scytonemin (OxScy), reduced scytonemin (ReScy) and carotenoids in order to reproduce the fine spectral shifts in the SVD spectra. See Supplementary Note 2 for detailed explanations. (a) Three spectral forms of OxScy. For the pigment-selected images as in Fig. 1d in the main text, the integrated intensity of only OxScy(high) and OxScy(low) between 1575 and 1625 cm^−1^ was used. (b) Two spectral forms of ReScy. The integrated intensity of both ReScy(high) and ReScy(low) between 1525 and 1580 cm^−1^ was used for the pigment-selective images as in Fig. 1e in the main text. (c) Four spectral forms of carotenoids. The integrated intensity of only carotenoids(high), carotenoids(middle), and carotenoids(low) between 1475 and 1525 cm^−1^ was used for the pigment-selective images as in Fig. 1b in the main text. (d) Spectra for phycobilins and chlorophyll. The integrated intensity between 1600 and 1650 cm^−1^ was used for the pigment-selective images of phycobilins as in Fig. 1c in the main text.

**Extended data Fig. 7.**
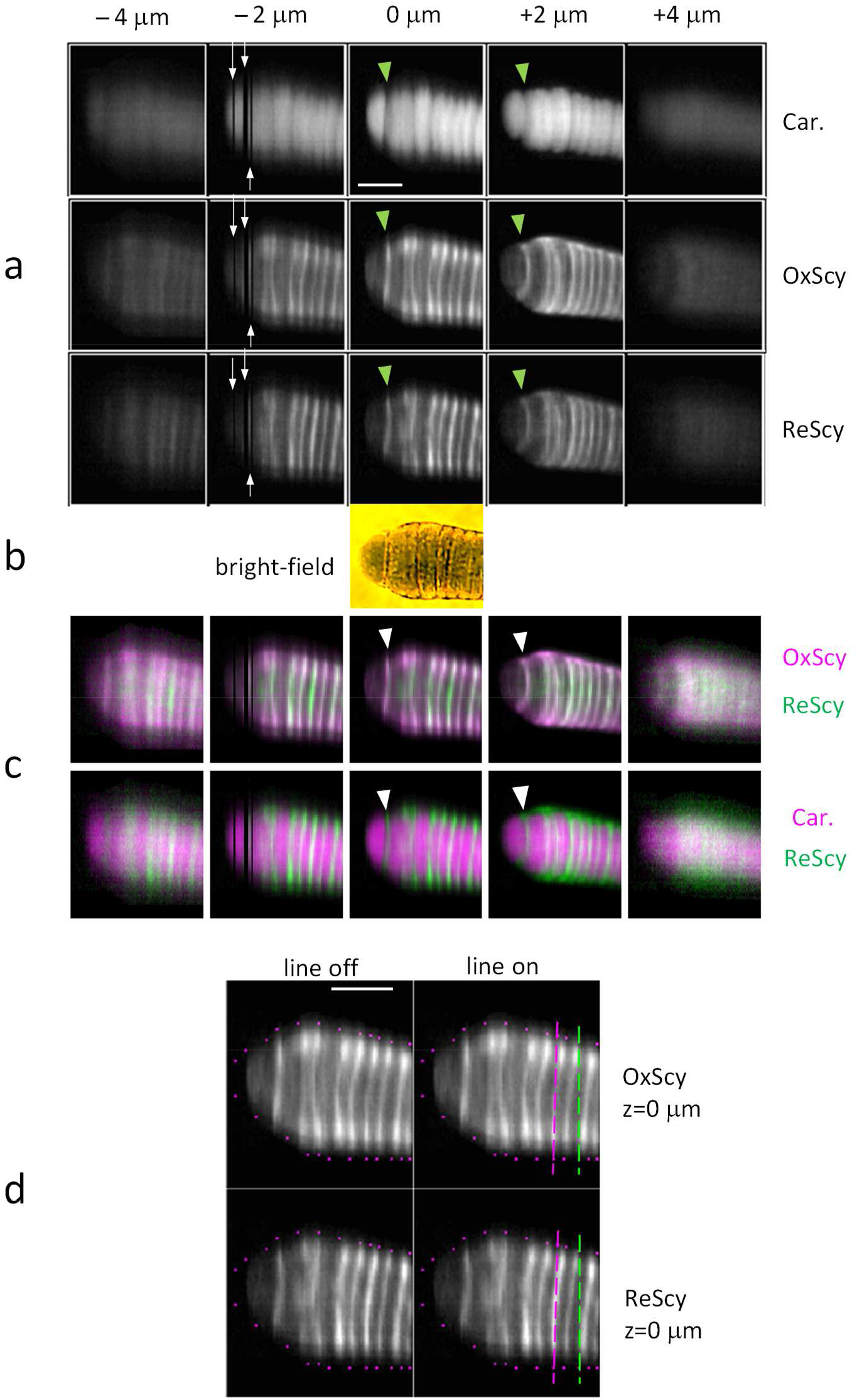
Pigment-selective Raman images at five depth positions (different z positions) of a filament that contained the reduced scytonemin without the UVA treatment. This filament was classified as [w/ ReDS, no UVA]. (a) Raman images of carotenoids (car.), oxidized scytonemin (OxScy) and reduced scytonemin (ReScy). Scale bar is 10 μm. The numbers at the top (−4 μm, - 2 μm, 0 μm, 2 μm, and 4 μm) show relative heights of the focus plane (z) in the cells. The upward or downward white arrows at the z position of - 2 μm indicate three regions for which Raman measurements of the cells were transiently interfered by unidentified contaminants. (b) Bright field image. (c) False-colored merged images for the pair of ReScy and OxScy (upper) and the one of ReScy and carotenoid (lower). The green arrow heads in (a) and white arrow heads in (c) indicate the position of heterocyst-vegetative cell junction. There are two points to be noted. First, ReScy is more internally located than OxScy as shown in the upper merged images in (c). See Supplementary Figs. 8 - 10 for filaments without substantial accumulatios of ReScy. Second, concentrated regions of carotenoids and ReScy are well separated as shown in the lower merged images in (c). (d) Enlarged images of OxScy and ReScy at 0 μm in (a) are marked by reference pixels and lines. The reference pixels (magenta dots) were selected for defining two contour lines along the long axis of the filament, which are close to the extracellular sheath. The green/magenta reference lines (broken line) were used to calculate the transverse cross section profiles in Fig. 4 of the main text, where the lines in magenta and green are approximately overlapping with the core of a cell junction and center of a cytoplasmic space, respectively.

**Extended data Fig. 8.**
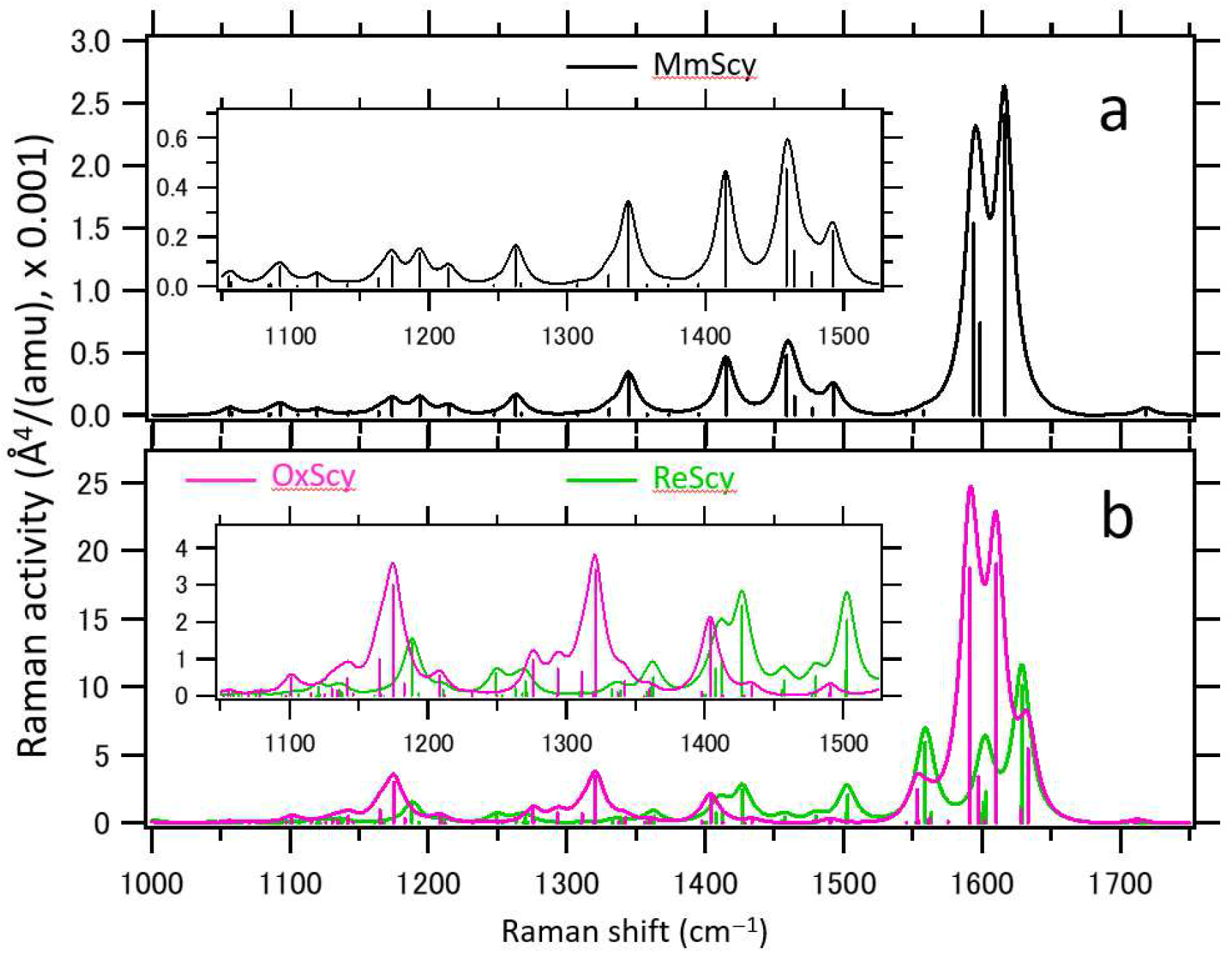
Quantum mechanical predictions of Raman spectra of scytonemins. (a) Monomer scytonemin (MmScy), (b) Oxidized scytonemin (OxScy, magenta) and reduced scytonemin (ReScy, green). The height of a single vertical bar represents Raman activity of one vibrational mode following a reference^38^, where Å is the angstrom unit and “amu” is the unified atomic mass unit. Integrated spectra of all vibrational modes of the specific molecules are shown by introducing a non-normalized Lorentzian broadening function with a full width at half maximum of 18 cm^−1^ and a peak value of 1. Accordingly, the unit of vertical axis is not applicable to the integrated spectra, but comparisons of the intensities between different plots of broadened molecular spectra remain meaningful. The inserts show magnified views of the graphs in the Raman shift range between 1050 and 1525 cm^−1^.

## Notes

### Competing Interest Statement

The authors have declared no competing interest.

## References

1. Fischer, W.W., Hemp, J. & Johnson, J.E. Evolution of oxygenic photosynthesis. Annual Review of Earth and Planetary Sciences 44, 647–683 (2016).

2. Holland, H.D. The oxygenation of the atmosphere and oceans. Philosophical Transactions of the Royal Society B: Biological Sciences 361, 903–915 (2006).

3. Shih, P.M. Early cyanobacteria and the innovation of microbial sunscreens. Mbio 10 (2019).

4. Gebauer, S., Grenfell, J.L., Lehmann, R. & Rauer, H. Effect of Geologically Constrained Environmental Parameters on the Atmosphere and Biosphere of Early Earth. ACS Earth and Space Chemistry 2, 1112–1136 (2018).

5. Takahashi, S. et al. The solar action spectrum of photosystem II damage. Plant Physiology 153, 988–993 (2010).

6. Nishiyama, Y. & Murata, N. Revised scheme for the mechanism of photoinhibition and its application to enhance the abiotic stress tolerance of the photosynthetic machinery. Applied microbiology and biotechnology 98, 8777–8796 (2014).

7. Garcia - Pichel, F. & Castenholz, R.W. Characterization and biological implications of scytonemin, a cyanobacterial sheath pigment 1. Journal of Phycology 27, 395–409 (1991).

8. Pathak, J. et al. Cyanobacterial Secondary Metabolite Scytonemin: A Potential Photoprotective and Pharmaceutical Compound. Proceedings of the National Academy of Sciences, India Section B: Biological Sciences 90, 467–481 (2020).

9. Garcia-Pichel, F. et al. Timing the evolutionary advent of cyanobacteria and the later great oxidation event using gene phylogenies of a sunscreen. Mbio 10, e00561–00519 (2019).

10. Colville, L. & Kranner, I. Desiccation tolerant plants as model systems to study redox regulation of protein thiols. Plant Growth Regulation 62, 241–255 (2010).

11. Matsui, K. et al. The cyanobacterial UV-absorbing pigment scytonemin displays radical-scavenging activity. The Journal of general and applied microbiology 58, 137–144 (2012).

12. Rastogi, R.P. & Incharoensakdi, A. Characterization of UV-screening compounds, mycosporine-like amino acids, and scytonemin in the cyanobacterium Lyngbya sp. CU2555. FEMS microbiology ecology 87, 244–256 (2014).

13. Schmitt, F.J. et al. Reactive oxygen species: Re-evaluation of generation, monitoring and role in stress-signaling in phototrophic organisms. Biochim. Biophys. Acta-Bioenerg. 1837, 835–848 (2014).

14. Pathak, J. et al. Cyanobacterial secondary metabolite scytonemin: a potential photoprotective and pharmaceutical compound. Proceedings of the National Academy of Sciences, India Section B: Biological Sciences, 1–15 (2019).

15. Klicki, K. et al. The widely conserved ebo cluster is involved in precursor transport to the periplasm during scytonemin synthesis in Nostoc punctiforme. MBio 9 (2018).

16. Squier, A.H., Hodgson, D.A. & Keely, B.J. A critical assessment of the analysis and distributions of scytonemin and related UV screening pigments in sediments. Organic Geochemistry 35, 1221–1228 (2004).

17. Itoh, T. et al. Reduced scytonemin isolated from Nostoc commune induces autophagic cell death in human T-lymphoid cell line Jurkat cells. Food and chemical toxicology 60, 76–82 (2013).

18. Edwards, H., Garcia-Pichel, F., Newton, E. & Wynn-Williams, D. Vibrational Raman spectroscopic study of scytonemin, the UV-protective cyanobacterial pigment. Spectrochimica Acta Part A: Molecular and Biomolecular Spectroscopy 56, 193–200 (2000).

19. Vítek, P. et al. Microbial colonization of halite from the hyper-arid Atacama Desert studied by Raman spectroscopy. Philosophical Transactions of the Royal Society A: Mathematical, Physical and Engineering Sciences 368, 3205–3221 (2010).

20. Vítek, P. et al. Distribution of scytonemin in endolithic microbial communities from halite crusts in the hyperarid zone of the Atacama Desert, Chile. FEMS microbiology ecology 90, 351–366 (2014).

21. Venckus, P., Paliulis, S., Kostkevičiene, J. & Dementjev, A. CARS microscopy of scytonemin in cyanobacteria Nostoc commune. Journal of Raman Spectroscopy 49, 1333–1338 (2018).

22. Nozue, S., Katayama, M., Terazima, M. & Kumazaki, S. Comparative study of thylakoid membranes in terminal heterocysts and vegetative cells from two cyanobacteria, Rivularia M-261 and Anabaena variabilis, by fluorescence and absorption spectral microscopy. Biochimica et Biophysica Acta (BBA)-Bioenergetics 1858, 742–749 (2017).

23. Tamamizu, K. & Kumazaki, S. Spectral microscopic imaging of heterocysts and vegetative cells in two filamentous cyanobacteria based on spontaneous Raman scattering and photoluminescence by 976 nm excitation. Biochimica et Biophysica Acta (BBA)-Bioenergetics 1860, 78–88 (2019).

24. Hirose, Y. & Katayama, M. Draft Genome Sequence of the Phototropic Cyanobacterium Rivularia sp. Strain IAM M-261. Microbiology Resource Announcements 10, e00790–00721 (2021).

25. Varnali, T. & Edwards, H.G. Reduced and oxidised scytonemin: theoretical protocol for Raman spectroscopic identification of potential key biomolecules for astrobiology. Spectrochimica Acta Part A: Molecular and Biomolecular Spectroscopy 117, 72–77 (2014).

26. Smith, E.M. et al. Survey of Scytonema (Cyanobacteria) and associated saxitoxins in the littoral zone of recreational lakes in Canterbury, New Zealand. Phycologia 51, 542–551 (2012).

27. El-Diasty, F. Coherent anti-Stokes Raman scattering: Spectroscopy and microscopy. Vibrational Spectroscopy 55, 1–37 (2011).

28. Gao, X., Jing, X., Liu, X.F. & Lindblad, P. Biotechnological Production of the Sunscreen Pigment Scytonemin in Cyanobacteria: Progress and Strategy. Marine Drugs 19 (2021).

29. De Philippis, R. & Vincenzini, M. Exocellular polysaccharides from cyanobacteria and their possible applications. FEMS Microbiology Reviews 22, 151–175 (1998).

30. Khayatan, B., Meeks, J.C. & Risser, D.D. Evidence that a modified type IV pilus - like system powers gliding motility and polysaccharide secretion in filamentous cyanobacteria. Molecular Microbiology 98, 1021–1036 (2015).

31. Wolk, C., Ernst, A. & Elhai, J. Heterocyst Metabolism and Development, in The Molecular Biology of Cyanobacteria. (ed. B. D.) 769–823 (Kluwer Academic Publisher, Netherland; 1994).

32. Pernil, R. & Schleiff, E. Metalloproteins in the Biology of Heterocysts. Life-Basel 9 (2019).

33. Sarasa-Buisan, C. et al. FurC (PerR) from Anabaena sp. PCC7120: a versatile transcriptional regulator engaged in the regulatory network of heterocyst development and nitrogen fixation. Environmental Microbiology.

34. Kaplan-Levy, R.N., Hadas, O., Summers, M.L., Rücker, J. & Sukenik, A. Akinetes: dormant cells of cyanobacteria. Dormancy and resistance in harsh environments, 5–27 (2010).

35. Natarajan, C., Prasanna, R., Gupta, V., Dureja, P. & Nain, L. Characterization of the fungicidal activity of Calothrix elenkinii using chemical methods and microscopy. Applied biochemistry and microbiology 48, 51–57 (2012).

36. Kozuch, S. & Martin, J.M. Spin - component - scaled double hybrids: an extensive search for the best fifth - rung functionals blending DFT and perturbation theory. Journal of Computational Chemistry 34, 2327–2344 (2013).

37. Weigend, F. & Ahlrichs, R. Balanced basis sets of split valence, triple zeta valence and quadruple zeta valence quality for H to Rn: Design and assessment of accuracy. Physical Chemistry Chemical Physics 7, 3297–3305 (2005).

38. Neugebauer, J., Reiher, M., Kind, C. & Hess, B.A. Quantum chemical calculation of vibrational spectra of large molecules - Raman and IR spectra for buckminsterfullerene. Journal of Computational Chemistry 23, 895–910 (2002).

39. Pinski, P. & Neese, F. Analytical gradient for the domain-based local pair natural orbital second order Moller-Plesset perturbation theory method (DLPNO-MP2). Journal of Chemical Physics 150 (2019).

40. Stoychev, G.L., Auer, A.A., Gauss, J. & Neese, F. DLPNO-MP2 second derivatives for the computation of polarizabilities and NMR shieldings. Journal of Chemical Physics 154 (2021).

41. Neese, F The ORCA program system. Wiley Interdisciplinary Reviews-Computational Molecular Science 2, 73–78 (2012).

42. Kesharwani, M.K., Brauer, B. & Martin, J.M.L. Frequency and Zero-Point Vibrational Energy Scale Factors for Double-Hybrid Density Functionals (and Other Selected Methods): Can Anharmonic Force Fields Be Avoided? Journal of Physical Chemistry A 119, 1701–1714 (2015).

43. Chan, B. & Radom, L. Frequency Scale Factors for Some Double-Hybrid Density Functional Theory Procedures: Accurate Thermochemical Components for High-Level Composite Protocols. Journal of Chemical Theory and Computation 12, 3774–3780 (2016).

44. Flores, E., Nieves-Morion, M. & Mullineaux, C.W. Cyanobacterial Septal Junctions: Properties and Regulation. Life-Basel 9 (2018).

45. Flores, E. & Herrero, A. Compartmentalized function through cell differentiation in filamentous cyanobacteria. Nature Reviews Microbiology 8, 39–50 (2010).

46. Hoiczyk, E. & Baumeister, W. The junctional pore complex, a prokaryotic secretion organelle, is the molecular motor underlying gliding motility in cyanobacteria. Current Biology 8, 1161–1168 (1998).

