## Supplementary information for "Sunscreen redox status in a multicellular cyanobacterium visualized by Raman scattering spectral microscope"

by

Kouto Tamamizu, Toshio Sakamoto, Yuki Kurashige, Shuho Nozue and Shigeichi Kumazaki*

### Table of contents

**Supplementary Fig. 1.**
Estimation of the spatial resolution of the line-scanning Raman spectral microscope.

**Supplementary Fig.2.**
A dark-sheathed *Rivularia* filament after the UVA treatment.

**Supplementary Fig. 3.** Correlation plot between the two indexes of heterocyst differentiation and accumulation of reduced scytonemin.

**Supplementary Fig. 4.** Histograms of pigment Raman signal ratios in the three filament groups.

**Supplementary Fig.5.**
Selected single-cell areas based on Raman imaging results.

**Supplementary Note 1.**
Details of singular value decompositions.

**Supplementary Fig. 6.**
Logarithmic plot of major singular values versus SVD component numbers for the representative filaments.

**Supplementary Note 2.**
Details of how we derived the component spectra of pigments.

**Supplementary Fig. 7.**
Maps of scores of ten major SVD components for a filament classified as [w/ ReDS, no UVA].

**Supplementary Fig.8.**
Images of a representative dark-sheathed *Rivularia* filament of the [w/ OxDS, +UVA] type.

**Supplementary Fig. 9.**

Images of a representative dark-sheathed Rivularia filament classified as [w/ OxDS, no UVA] that had OxScy mainly at the extracellular sheaths.

**Supplementary Fig. 10.**

Images of a dark-sheathed Rivularia filament classified as [w/ OxDS, no UVA] that had OxScy in the regions close to cell junctions as well as at the extracellular sheath.

**Supplementary Fig. 11.**

Pigment-Raman signal profiles along longitudinal axes of the filaments of the [w/ ReDS, no UVA] type.

**Supplementary Fig. 12.**

Pigment-Raman signal profiles along longitudinal axes of the filaments of the [w/ OxDS, +UVA] type.

**Supplementary Fig. 13.**

Theoretical visualization of one vibrational mode that is mainly associated with the carbon-carbon bond (C1-C22) of the oxidized or reduced scytonemin (See Extended data Fig. 1 for the atomic numbering).

**Supplementary Fig. 14.**

Theoretical visualization of three vibrational modes of the monomeric scytonemin (MmScy).

**Supplementary Fig. 15.**

Theoretical visualization of five vibrational modes of the reduced scytonemin (ReScy) that are candidates for the experimentally observed Raman band at 1365 – 1366 $cm^{-1}$ in the Rivularia filament and in the solid-state reduced scytonemin.

**Supplementary Fig. 16.** The ratio profiles of phycobilins/carotenoids along longitudinal filament axes.

**Supplementary Fig. 17**. Picture of thick accumulations of *Rivularia* filaments on agar media in glass bottles.

**Supplementary Fig. 18.** Schematic drawing of a preculture of *Rivularia* filaments before microscopic observations.

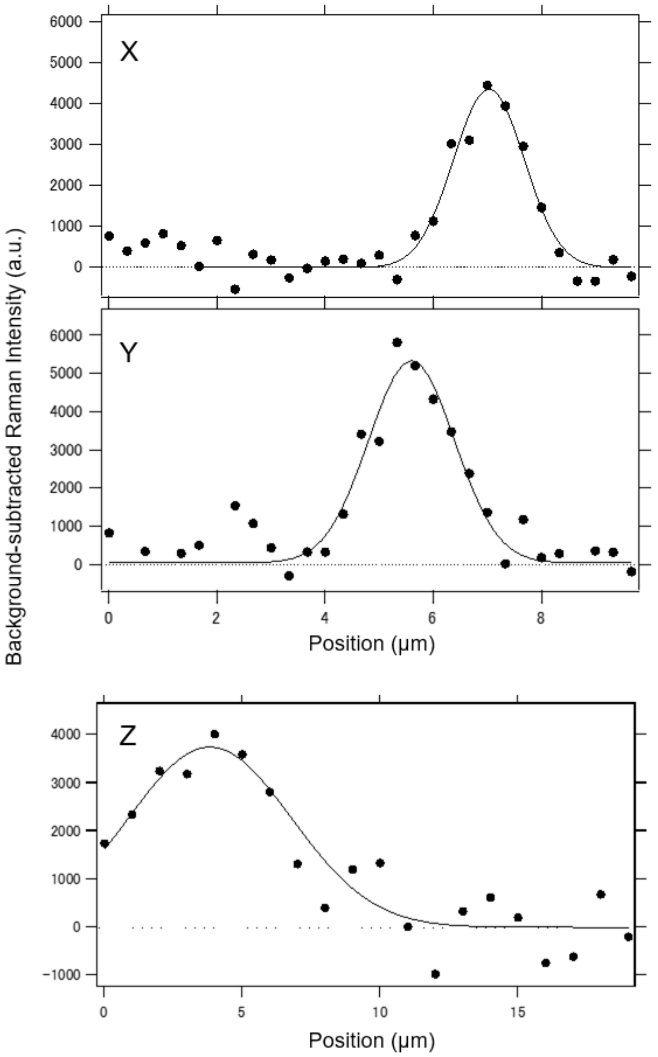


**Supplementary Fig. 1**. Estimation of the spatial resolution of the line-scanning Raman spectral microscope. Experimental Raman signal profiles integrated between 995 and 1011 $cm^{-1}$ of polystyrene beads (Funakoshi, no.19814) with a diameter of 2.0 μm are shown by closed circles. The data along X, Y, Z axes were best fitted by Gaussian functions (solid line) with full width at half maxima (FWHM) of 1.37, 1.72, and 6.09 μm, respectively. We carried out numerical simulations to make convolutions of three dimensional (3D) Gaussians (point-spread functions) of various FWHMs with a 3D sphere with a diameter of 2.0 μm fitted to simple 3D Gaussian functions.  Based on these calculations, the FWHM of the point-spread function of our Raman microscope were estimated to be 0.51, 1.1 and 6.0 mm, respectively. It is to be noted that the estimated FWHM of the point spread function along the X axis was near the diffraction limit given by 0.61×λ/NA (0.61×1.064/1.27= about 0.51 μm).

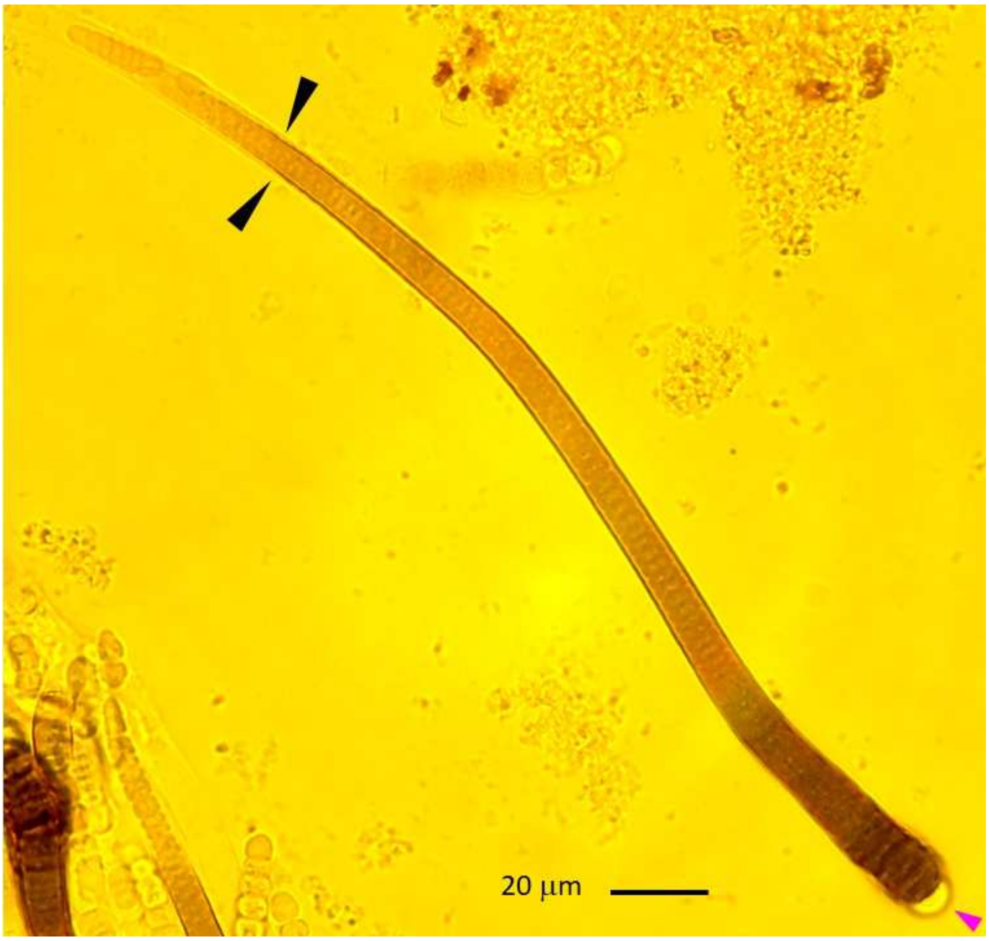


**Supplementary Fig. 2.**

**A dark-sheathed *Rivularia* filament after the UVA treatment**. This filament was not included as a target of Raman microscopy, but shown here as an example of filaments having dark-sheaths, because of the better contrast and similar coverage of the dark sheaths over the whole filament in this image in comparison with those recorded together with the Raman microscopy. The magenta arrow head indicates the terminal heterocyst at the basal end of the filament, and the two black ones indicate dark extracellular sheaths close to the tapered end (tip end) of the filament. The scale bar indicates 20 μm. It is to be noted that dark-sheaths of *Rivularia* filaments without the UVA treatment were often limited to the basal region, as shown in Fig. 1 in the main text.

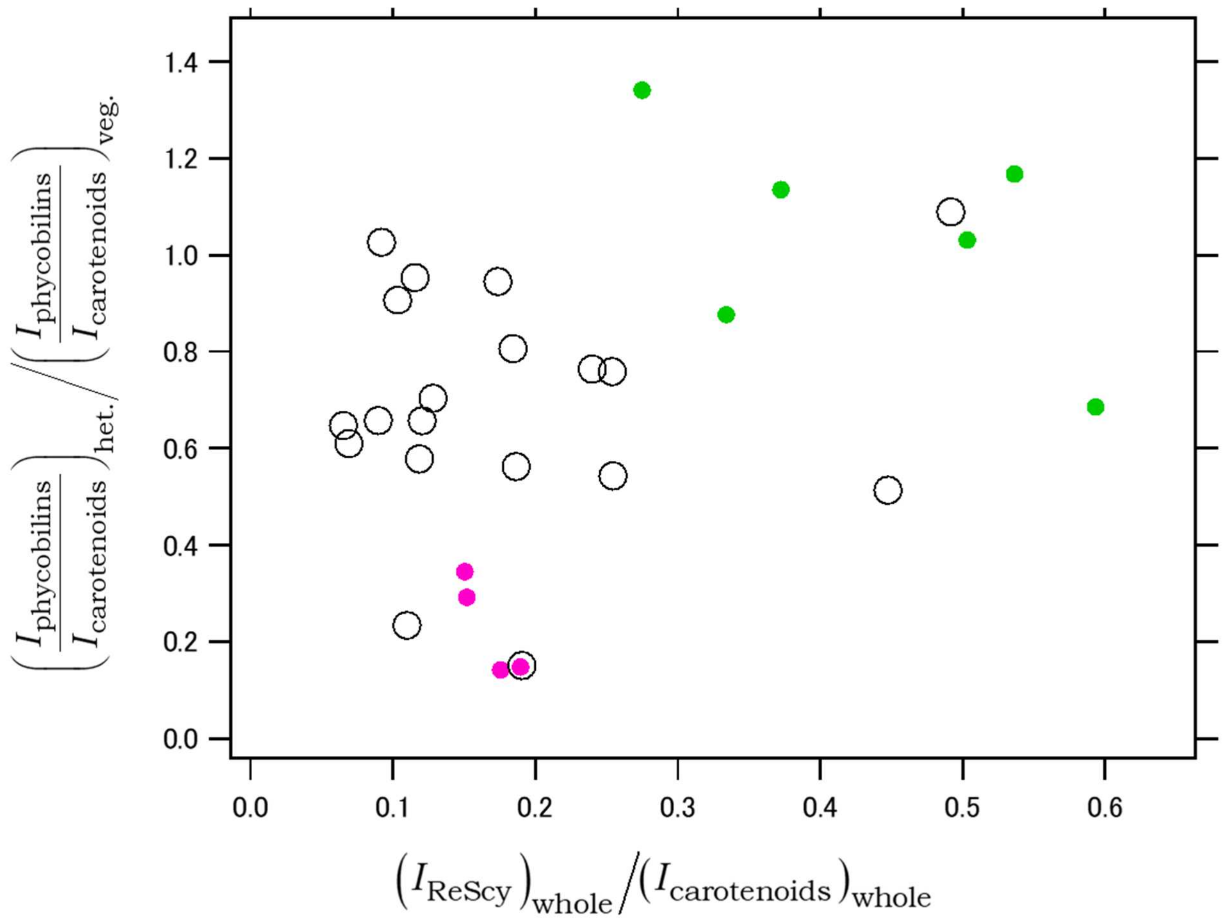


**Supplementary Fig. 3.** Correlation plot between the two indexes of heterocyst differentiation and accumulation of reduced scytonemin. The horizontal and vertical axes represent indexes of accumulation of reduced scytonemin (ReScy) and heterocyst differentiation, respectively, as detailed in the legend of Table 1 in the main text. Open black circles, closed magenta dots, and closed green dots represent single filaments classified as [w/ OxDS, no UVA], [w/ OxDS, +UVA] and [w/ ReDS, no UVA].

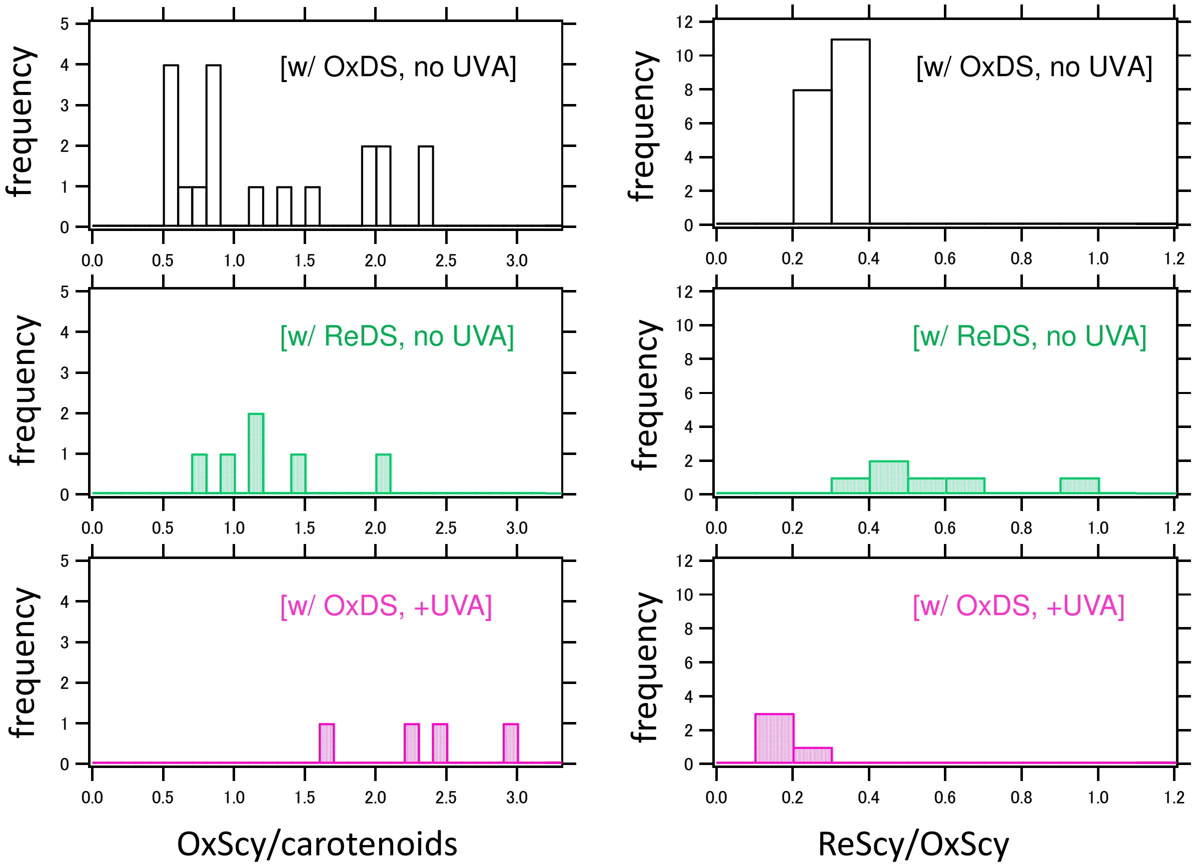


Supplementary Fig. 4. Histograms of pigment Raman signal ratios in the three filament groups. The definitions of the ratios are defined in the legend of Extended data Table 1.

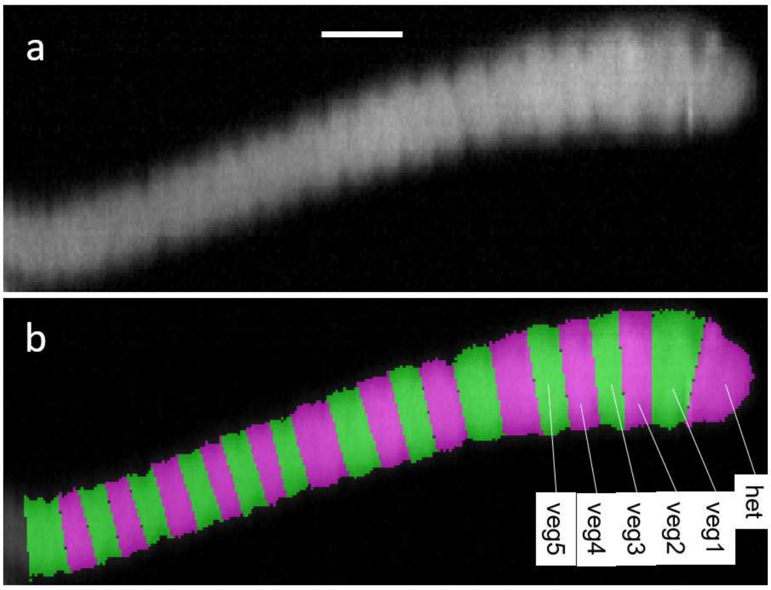


**Supplementary Fig. 5. Selected single-cell areas based on Raman imaging results.** (a) Image of approximate carotenoid-rich Raman signals between 1141 and 1179 cm$^{-1}$ of the dark-sheathed filament classified as [w/ ReDS, no UVA] that is identical to the one in Fig. 1 in the main text. The scale bar (10 μm) is also applicable to (b). (b) Map of selected areas of individual cells of the filament in (a) for spectral analysis. Magenta and green regions alternately show selected regions of individual cells.

**Supplementary Note 1. Details of singular value decompositions.**

Raman microscopic imaging produced Raman spectra at many spatial points, which was expressed as a matrix as follows.

$$\begin{aligned} &R(x_l, \lambda_i) \\ &l = 1, 2, ......, n-1, n \\ &i = 1, 2, ......, m-1, m \end{aligned} , \qquad \text{(eq. 1)}$$

where ($x_l$, $\lambda_i$) denote a pair of certain spatial position and wavelength (vibrational frequency) distinguished from the others by numerical subscripts of *l* and *i*, and *n* and *m* denote the total numbers of spatial points and wavelength points (vibrational frequencies), respectively. The singular value decomposition (SVD) function of Mathematica (Wolfram Research) was applied to the matrix *R*, which yielded the following factorized form.

$$R = U \Sigma V^T , \quad \text{(eq.2)}$$

where $U$ was an $n \times n$ real orthogonal matrix, $\Sigma$ was a $n \times m$ rectangular diagonal matrix with non-negative real numbers (singular values) on the diagonal, and $V^T$ was the transposed matrix of a matrix $V$ that was a $m \times m$ real orthogonal matrix. Representative examples of the major diagonal values (called singular values) of the three filaments of the different types are plotted in Supplementary Fig. 6. By selecting relatively major diagonal elements, the matrix $R$ was approximated as follows.

$$R = U\Sigma V^T$$
$$\approx \begin{pmatrix} u_1(x_1) & u_2(x_1) & u_3(x_1) & 0 & \cdots \\ u_1(x_2) & u_2(x_2) & u_3(x_2) & 0 & \cdots \\ \vdots & \vdots & \vdots & \vdots & \\ u_1(x_{n-1}) & u_2(x_{n-1}) & u_3(x_{n-1}) & 0 & \cdots \\ u_1(x_n) & u_2(x_n) & u_3(x_n) & 0 & \cdots \end{pmatrix} \begin{pmatrix} \sigma_{11} & 0 & 0 & 0 & \cdots \\ 0 & \sigma_{22} & 0 & 0 & \cdots \\ 0 & 0 & \sigma_{33} & 0 & \cdots \\ 0 & 0 & 0 & 0 & \cdots \\ \vdots & \vdots & \vdots & \vdots & \ddots \end{pmatrix} \begin{pmatrix} f_1(\lambda_1) & f_1(\lambda_2) & \cdots & & f_1(\lambda_m) \\ f_2(\lambda_1) & f_2(\lambda_2) & \cdots & & f_2(\lambda_m) \\ f_3(\lambda_1) & f_3(\lambda_2) & \cdots & & f_3(\lambda_m) \\ 0 & 0 & \cdots & 0 & 0 \\ \vdots & \vdots & & \vdots & \vdots \end{pmatrix}$$

(eq. 3)

where the above equation (eq.3) is an example in the case of selecting only three major singular values. Examples of the set of values: $f_1(\lambda_i)$, $f_2(\lambda_i)$, $f_3(\lambda_i)$, …., $f_{10}(\lambda_i)$, where $\lambda_i$ spans all the measured wavelength (vibrational frequency) range ($i$ =1,2,…., $m$) in the case of selecting ten major singular values, are shown in the Extended data Figs. 3 and 4. Those spectra are called "SVD spectra" in this work, and they possibly show both positive and negative values. The SVD spectra in general need to be expressed by sum and/or difference of multiple spectra that consist of non-negative values attributable to different molecules (spectral decomposition). Examples of the set of values: $u_1(x_l)$, $u_2(x_l)$, $u_3(x_l)$, …., $u_{10}(x_l)$, where $x_l$ spans all the considered spatial points (pixels, $l$=1,2,….,$n$) in the case of selecting ten major singular values, are shown in the Supplementary Fig. 7. Those maps are called "maps of scores" in this work, and they also possibly show both positive and negative values. Spectral decompositions of the SVD spectra (e.g., Extended data Figs. 3 - 5) yielded conversion coefficients between the SVD spectra and the molecular component spectra (Extended data Fig. 6) that were used to convert the maps of scores into concentration maps of pigments (e.g., Fig. 1, Extended data Fig. 7, Supplementary Figs. 8 - 10)

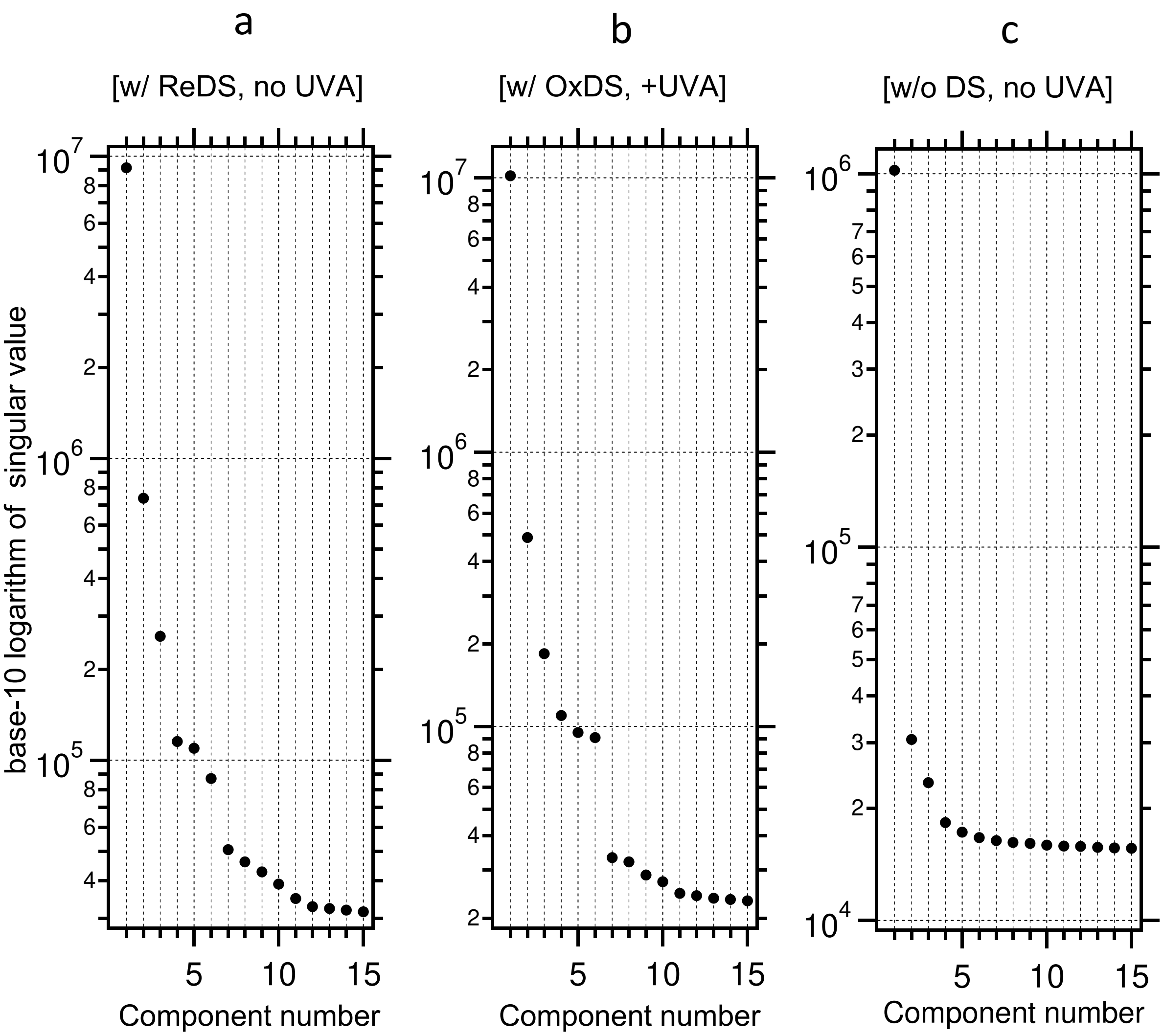


**Supplementary Fig. 6. Logarithmic plot of major singular values versus SVD component numbers for the representative filaments.** (a) dark-sheathed filament having ReScy without the UVA treatment, which was classified as [w/ ReDS, no UVA], (b) dark-sheathed filament after the UVA treatment that was classified as [w/ OxDS, +UVA], (c) filament without dark sheaths, classified as [w/o DS, no UVA]. The accumulations of OxScy and/or ReScy in the filament types of [w/ ReDS, no UVA] and [w/ OxDS, +UVA] necessitated about 10 SVD spectra, while only about 4 SVD spectra were necessary in the filament type of [w/o, DS, no UVA] (Extended data Figs. 3 - 5).

### Supplementary Note 2. Details of how we derived the component spectra of pigments.

The number of significant SVD spectra (Supplementary note 1) was smallest in the case of the [w/o DS, no UVA] (Extended Fig. 5, Supplementary Fig. 6c). We thus started preparation of component spectra based on the SVD spectra in the Extended data Fig. 5 and the other SVD spectra of the same filament type (all three filaments classified as [w/o DS, no UVA]). First, several Raman-shift regions (frequency regions) were assumed to be free from Raman signals and that they show only some autoluminescence signals. In the case of the second SVD of the Extended data Fig. 5 (SVD2), for example, regions between the following lower and upper limits were assumed to be Raman-free regions: (743, 901), (1090, 1095), (1690, 1775) in cm$^{-1}$. Different sets of Raman-free regions were selected for individual SVD spectra. The whole spectra of autoluminescence amplitudes contributing to the SVD spectra were assumed to be given by copies of SVD spectral amplitudes in the Raman-free regions and straight lines connecting the neighboring Raman-free regions. The fitting curves in the Extended data Figs 3 - 5 were given by sums of such autoluminescence spectral amplitudes and Raman spectra of pigments.

The first SVD spectrum of one *Rivularia* filament of the type of [w/o DS, no UVA] (SVD1, Extended data Fig. 5) showed three negative-going Raman bands characteristic to carotenoids at 1006, 1159, and 1521 cm$^{-1}$, as reported in our previous work[1]. The second SVD spectrum (SVD2) showed two positive-going Raman bands at 1159 and 1522 cm$^{-1}$ that are nearly indistinguishable from those of SVD1. The peak position of the Raman bands at around 1006 cm$^{-1}$ of the SVD2 spectrum was not clearly determined, but corresponding well to that of SVD1. The fourth SVD spectrum (SVD4) showed sinusoidal shapes in narrowly bounded regions around the three carotenoid Raman bands with unified positive/negative alternations. The other two sets of the SVD spectra of the other two *Rivularia* filaments of the same [w/o DS, no UVA] group also included a SVD spectrum showing the same spectral features. It was also reported that center positions of the two Raman bands of carotenoids at around 1160 and 1520 cm$^{-1}$ are correlated among many types of carotenoids[2]. Thus, it is likely that the spectral features of the SVD4 indicates at least two types of carotenoid molecules with different spatial distributions. These considerations led us to assume three carotenoid spectra that were able to reproduce both upward and downward frequency shifts from central frequencies (Extended data Fig. 6c). In each filament, the central frequencies of the carotenoid Raman bands were determined through curve fittings of a selected SVD spectrum that show carotenoid signals as purely as possible and has a singular value as large as possible. The fourth carotenoid spectrum was also assumed (Extended data Fig. 6c), which will be explained below. Based on Raman spectra in our previous work and references, all Raman bands including those of carotenoids were given by Lorentzian or Gaussian functions of which peak positions and bandwidths were finely adjusted through curve fittings. More analysis on the presence of multiple carotenoids will be addressed in our future publications, and we here just consider sums of signals of all carotenoids (including only three major spectral types and excluding the fourth minor carotenoid that is described below) in the main text.

In addition to carotenoids, phycobilins and chlorophylls are major photosynthetic pigments in cyanobacteria and they are also present in *Rivularia* M-261 cells[1, 3]. Phycobilin Raman bands were clearly visible mainly as two negative-going signals centered at about 1374 and 1629 cm$^{-1}$ in the third SVD spectra (SVD3, Extended data Fig. 5) as well as those at 1371 and 1639 cm$^{-1}$ in the cellular Raman spectra of vegetative cells of the filament type of [w/o DS, no UVA] (Fig. 2b). Based on these signals and several literatures, we constructed the Raman spectrum of phycobilins (Extended data Fig. 6d). Although Raman signals of chlorophylls were recognizable only in the first SVD spectrum of the filament classified as [w/o DS, no UVA] (for example, one band at about 1325 cm$^{-1}$ in SVD1, Extended data Fig. 5a), Raman spectrum of chlorophyll was constructed and included in the curve fittings by using several references and our own previous work[1].

The above-mentioned considerations yielded Raman spectra of carotenoids, phycobilins and chlorophylls. These spectra were also used in the curve fittings of the *Rivularia* filaments of the other types. Raman signals of oxidized scytonemin (OxScy) at around 1175 and 1595 $cm^{-1}$ were visible at least in the first, third, fourth, fifth, sixth and seventh SVD spectra of the filaments of the [w/ OxDS, +UVA] type (Extended data Fig. 4). We found it difficult to reproduce all SVD spectra by a single OxScy spectrum. We thus introduced two spectra of OxScy with slight frequency shifts (OxScy(high) and OxScy(low) in Extended data Fig. 6a). We also introduced a third OxScy Raman spectrum (Scy(minor) in Extended data Fig. 6a) that will be described below.

In addition to Raman signals of carotenoid, phycobilins, chlorophylls and OxScy, Raman signals of ReScy were needed to reproduce the negative-going signals around 1070, 1367 and 1550 $cm^{-1}$ in the fifth SVD spectrum of the filament classified as [w/ ReDS, no UVA] (SVD5, Extended data Fig. 3). These Raman bands also seem to be appearing as positive-going signals in the fourth SVD spectrum (SVD4) and negative-going signals in the third SVD spectrum (SVD3, Extended data Fig. 3). Based mainly on these spectral features, Raman spectrum of ReScy was prepared (Extended data Fig. 6b). However, it was again difficult for the single Raman spectrum of ReScy to reproduce the complicated spectral shapes around 1550 $cm^{-1}$ of some SVD spectra (especially SVD7 and SVD8 spectra in the Extended data Fig. 3). We thus tentatively assumed two Raman spectra with a slight shift in the peak position of the band around 1550 $cm^{-1}$ ((ReScy(high) and ReScy(low) in Extended data Fig. 6b).

Although sum of autoluminescence spectra, the three spectra of carotenoids (high, middle, low), one of phycobilin, one of chlorophyll, two spectra of OxScy (high, low), and two spectra of ReScy (high, low) with varied weights were reasonably successful to reproduce most SVD spectra (Extended data Fig. 3 - 6), there remained some discrepancies. We tentatively introduced a third scytonemin Raman spectrum and fourth carotenoid Raman spectrum to further improve the curve fittings (Scy(minor) and carotenoid(minor) as in Extended data Fig. 6). These two spectral components were included in the fitted curves shown in the fittings of SVD spectra (Extended data Fig. 3 - 5), but they were not considered in the calculation of maps and profiles of carotenoids and OxScy in the other figures (Fig. 1,3,4, Table 1, Extended data Fig. 7, Supplementary Figs. 8 - 12,16). It should be noted here that the spectral features of the minor carotenoid (carotenoid (minor) in Extended data Fig. 6c) in this study bears a certain similarity to those of extracellular polymeric substances around cyanobacteria reported in a previous study [4]. We also found a weak positive correlation in the spatial distribution between the "minor carotenoid" and scytonemins (data not shown). It is thus possible either that the "minor carotenoid" is a special carotenoid not localized around thylakoid membranes or that the "minor carotenoid" spectrum indicates different substances constituting EPS sheaths.

The scripts used to perform the above-mentioned curve fittings of SVD spectra by sum of pigment Raman spectra are provided as a supplemental file with this paper.

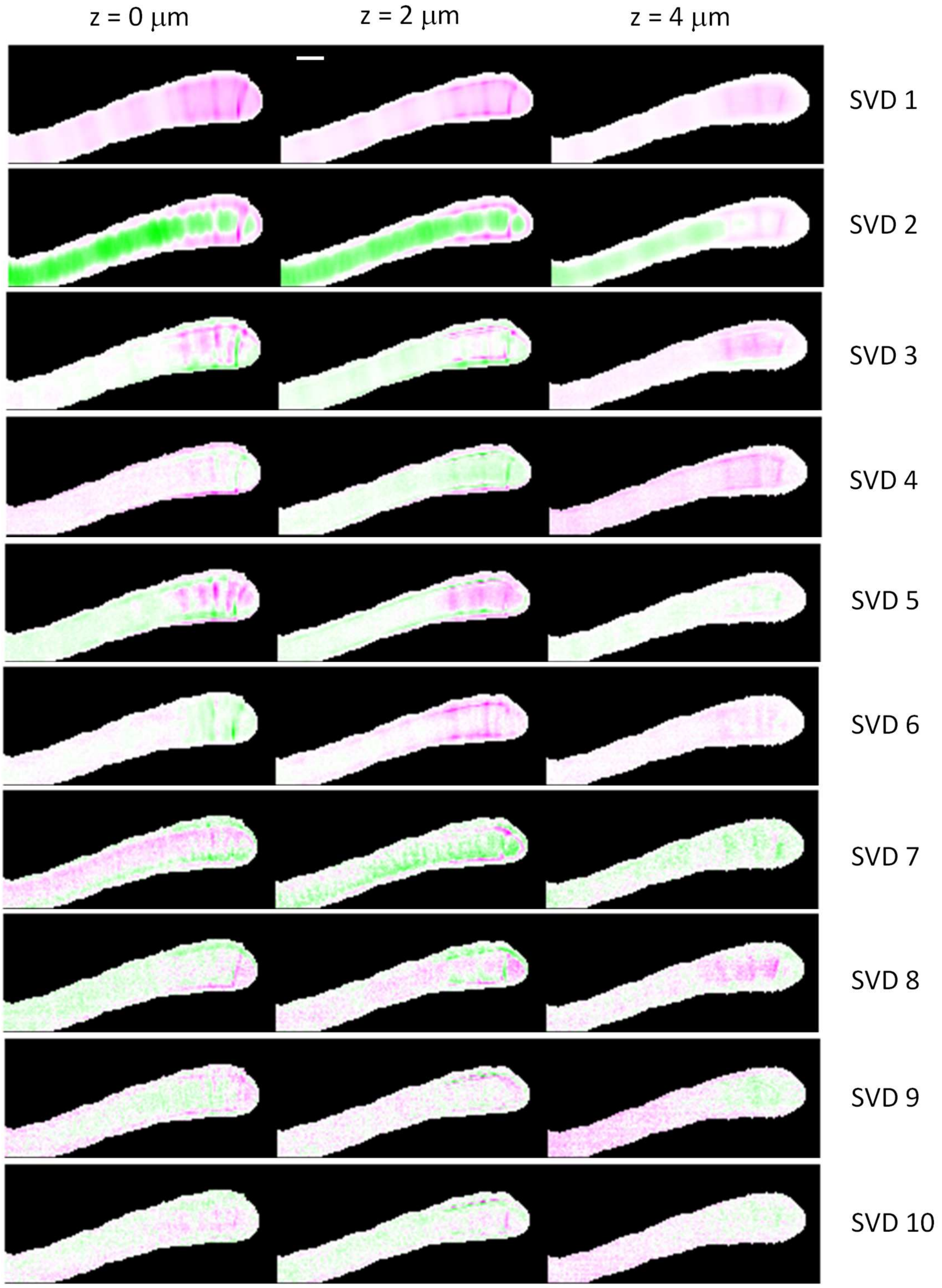


**Supplementary Fig. 7. Maps of scores of ten major SVD components for a filament classified as [w/ ReDS, no UVA].** The target filament was the same as that shown in Fig.1 of the main text. Positive and negative score values are shown by green and magenta, respectively. The whiter the color of a pixel, the smaller the absolute value in that pixel. Black regions were excluded from the SVD analysis. Scale bar is 10 μm. See Supplementary note 1 for the definition of these maps.

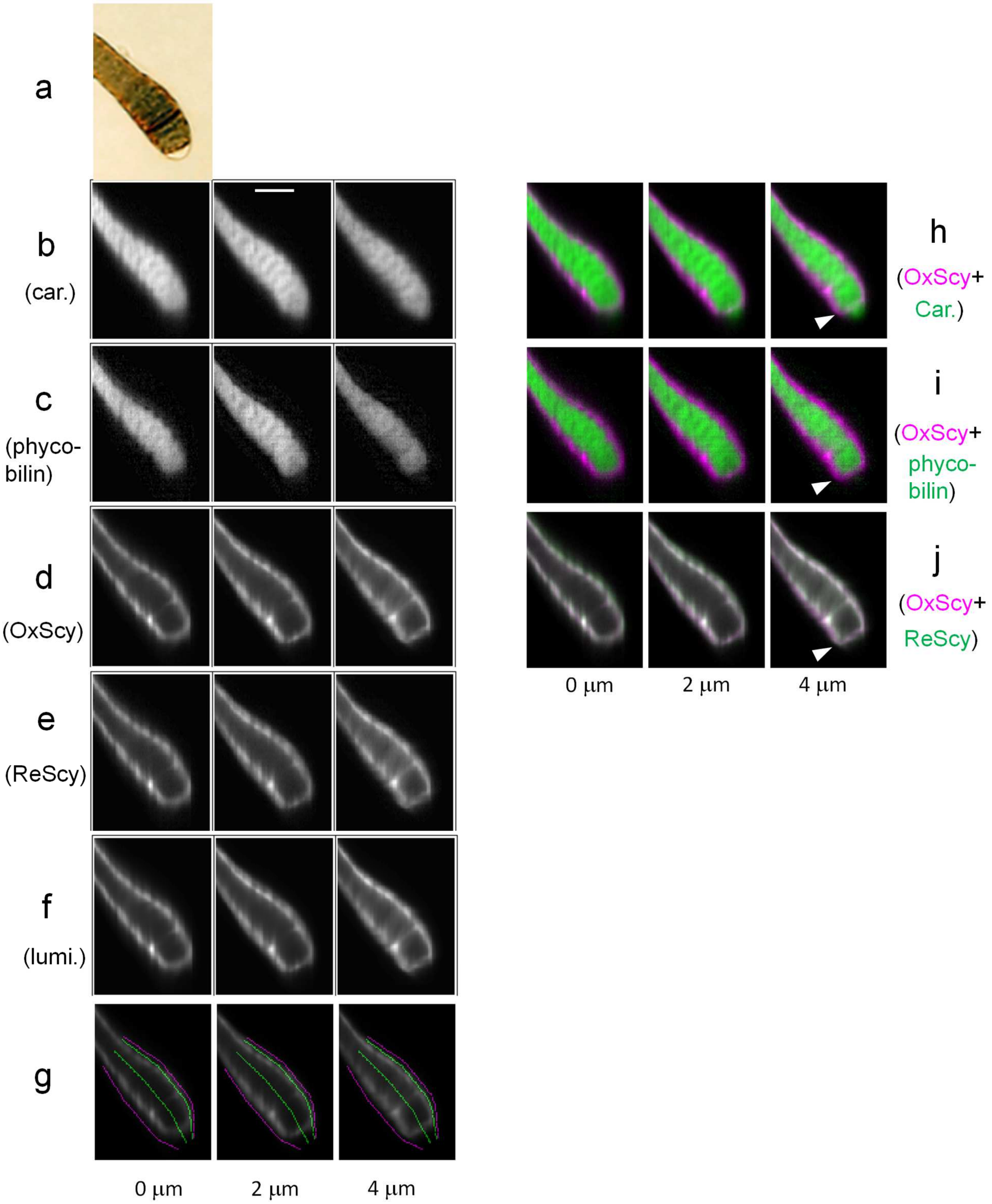


**Supplementary Fig. 8. Images of a representative dark-sheathed *Rivularia* filament of the [w/ OxDS, +UVA] type.** This figure was prepared in the same manner as Fig. 1 in the main text. The scale bar indicates 10 µm. Although the inclusion of the ReScy in the spectral decomposition of the filaments treated with UVA was positively supported neither by the SVD spectra (Extended Fig. 4) nor by the raw cellular spectra (Fig. 2 in the main text), ReScy was included in the spectral decomposition, and the resulting images of the ReScy component are shown here (as in (e)) for a fair comparison of the filaments of different types. There are three points to be noted here. First, the distribution of ReScy is very similar to that of

OxScy (See (j) for the false-colored merged image of OxScy and ReScy), suggesting that accumulation of ReScy is negligible. Second, there are no substantial accumulations of OxScy in the central regions of the vegetative-vegetative cell junctions at the relative heights (z) of 0 and 2 μm. Third, the distribution of carotenoids extends beyond the heterocyst-vegetative cell junction (indicated by white arrow head in (h - j)), while the one of phycobilins seems to be primarily limited to vegetative cells.

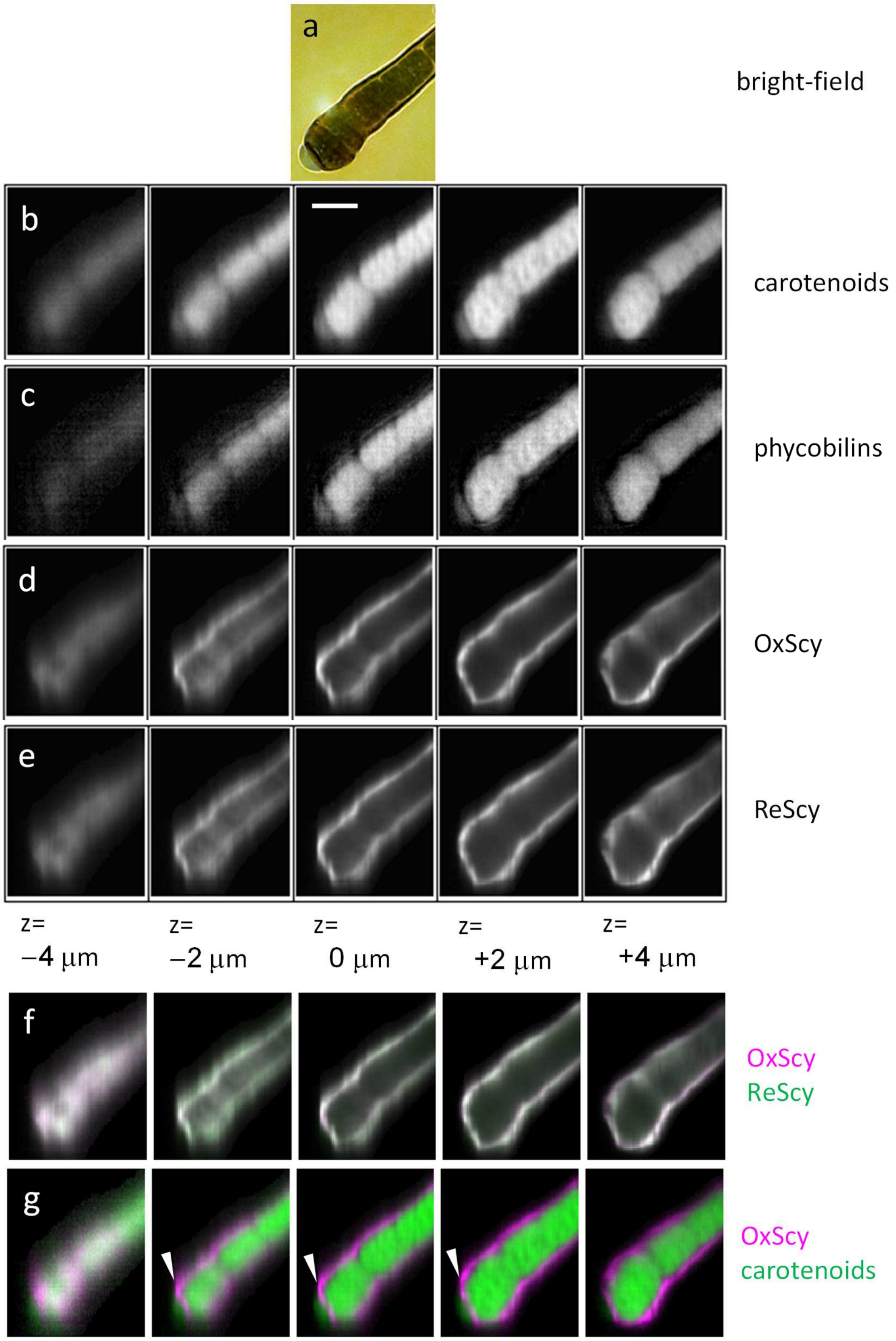


**Supplementary Fig. 9. Images of a representative dark-sheathed *Rivularia* filament classified as [w/ OxDS, no UVA] that had OxScy mainly at the extracellular sheaths.** This figure was prepared in the same manner as Fig. 1 in the main text. The scale bar indicates 10 μm. Although the inclusion of the ReScy in the spectral decomposition of this filaments was positively supported neither by the SVD spectra nor by the raw cellular spectra, ReScy was included in the spectral decomposition, and the resulting images of the ReScy component are shown here for a fair comparison of different filaments. There are four points to be noted here. First, the distribution of ReScy is very similar to that of OxScy (See (f) for false-colored merged image of OxScy and ReScy), suggesting that accumulation of ReScy is negligible. Second, there is no substantial accumulations of OxScy in the central regions of the vegetative-vegetative cell junctions at the relative heights (z) of 0 and 2 µm. Third, the distribution of carotenoids extends beyond the heterocyst-vegetative cell junction (indicated by white arrow heads in (g)). These features are very similar to those in the [w/ OxDS, +UVA] (Supplementary Fig. 8). Fourth, the heterocyst-vegetative cell junction is characterized by a local minimum of the concentration of carotenoids and local maximum of OxScy, which is indicated by the clearly magenta line between the heterocyst and connected vegetative cell that are colored green (in (g)).

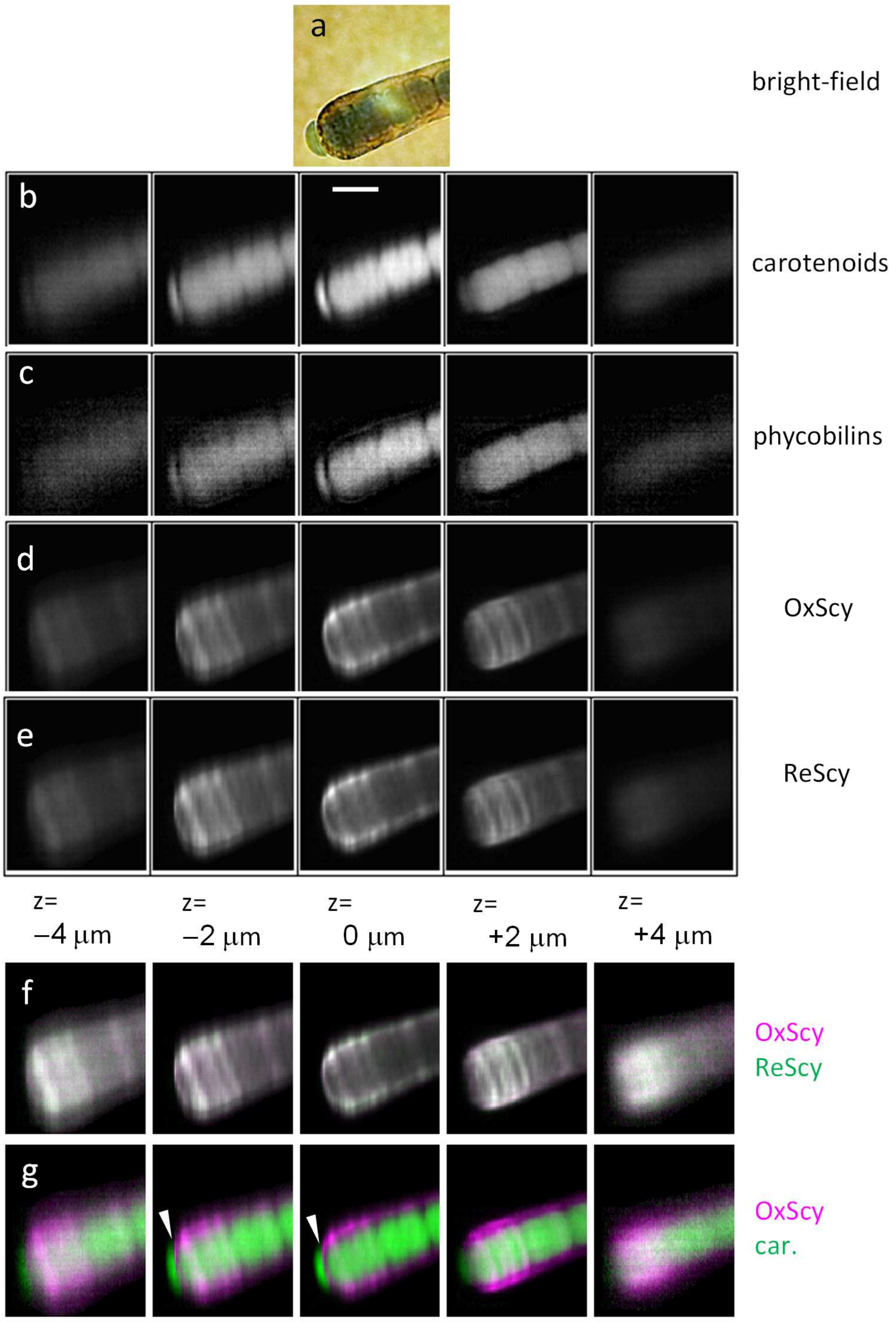


**Supplementary Fig. 10. Images of a dark-sheathed *Rivularia* filament classified as [w/ OxDS, no UVA] that had OxScy in the regions close to core of cell junctions as well as at the extracellular sheath.** This figure was prepared in the same manner as those described in the caption of Fig. 1 in the main text. The scale bar indicates 10 μm. Although the inclusion of the ReScy in the spectral decomposition of this filament was positively supported neither by the SVD spectra nor by the raw cellular spectra, ReScy was included in the spectral decomposition, and the resulting images of the ReScy component are shown here for a fair comparison of different filaments. There are four points to be noted here. First, the distribution of ReScy is very similar to that of OxScy (See (f) for false-colored merged image of OxScy and ReScy), suggesting that accumulation of ReScy is negligible. Second, in contrast to the filament in Supplementary Fig. 9, there are substantial accumulations of OxScy in the central regions of the vegetative-vegetative cell junctions even at central relative heights (z). Third, the distribution of carotenoids extends beyond the heterocyst-vegetative cell junction (indicated by white arrow heads in (g)). These features are very similar to those in the [w/ OxDS, +UVA] (Supplementary Fig. 8). Fourth, the heterocyst-vegetative cell junction is characterized by a local minimum of the concentration of carotenoids and local maximum of that of OxScy, which is indicated by the clearly magenta line between the heterocyst and connected vegetative cell that are colored green (in (g)).

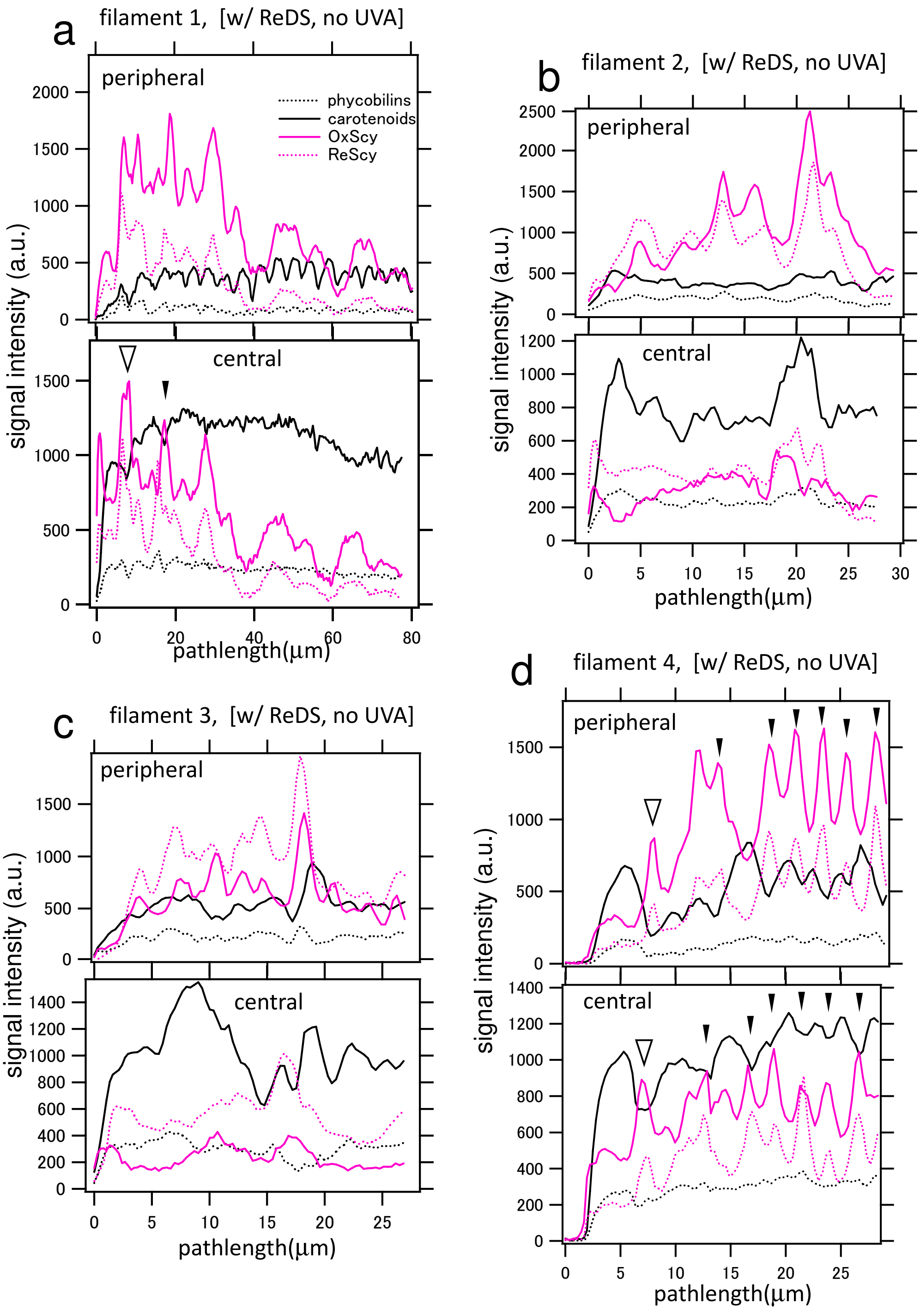


**Supplementary Fig. 11. Pigment-Raman signal profiles along longitudinal axes of the filaments containing the reduced scytonemin ([w/ ReDS, no UVA] type)**. The four individual panels (a - d) show Raman-signal profiles of pigments along the longitudinal axes of four *Rivularia* filaments (designated by the numbers from 1 to 4). The abscissa shows the pathlength from the basal end (heterocystous terminal) of the filament. Some of the profiles of the filament 1 are also shown in Fig. 3a of the main text. In each panel, the upper graph shows profiles along a line that is close to the extracellular sheath (peripheral line). The lower panel shows profiles along a line that is central and mainly in cytoplasmic regions (central line). The selections of the central and peripheral lines are analogous to the green lines shown in Fig.1g of the main text. In all plots, the intensities were not normalized so that their magnitudes are to be compared within the same panel. In (a) and (d), the open and closed black arrow heads indicate some relatively well resolved positions of heterocyst-vegetative and vegetative-vegetative cell junctions, respectively, at which signals of carotenoids show local minima while those of scytonemins show local maxima.

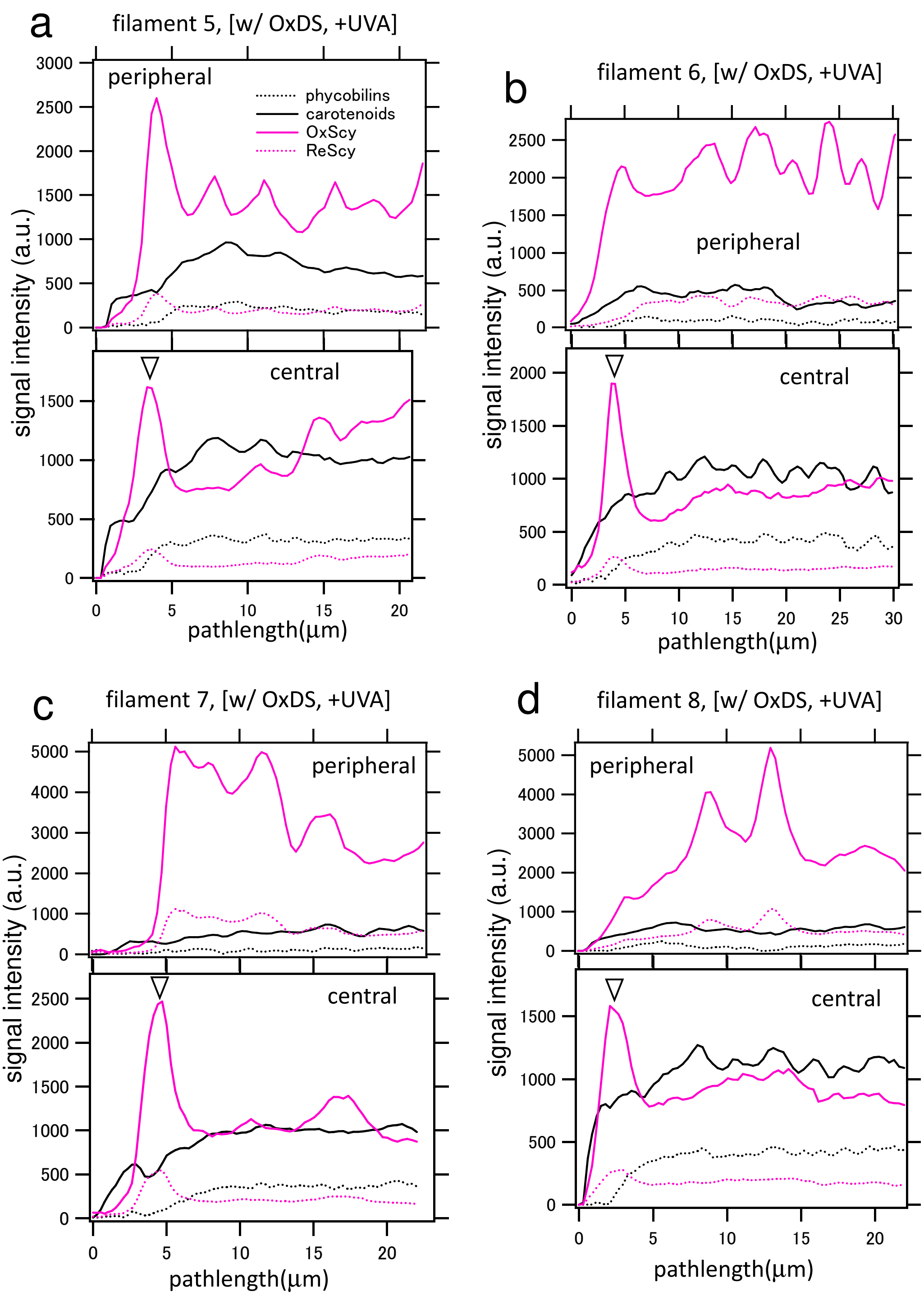


Supplementary Fig. 12. **Pigment-Raman signal profiles along longitudinal axes of the *Rivularia* filaments treated with the UVA illumination ([w/ OxDS, +UVA] type)**. The four individual panels (a - d) show Raman-signal profiles of pigments along longitudinal axes of four filaments (designated by the numbers from 5 to 8). Some of the profiles of the filament 6 (in the b panel) are shown in Fig. 3b of the main text. All plots, graphs and panels were prepared in the same manner as in Supplementary Fig. 11. The position of heterocyst-vegetative cell junctions is indicated by open black downward arrow heads for the central longitudinal axes. There are two noteworthy points in these data. (i) The ratio of phycobilins/carotenoids becomes significantly decreased in the heterocyst region (around 0 – 4 μm) than in the vegetative cell regions (>5 µm), the ratio of which is shown in Supplementary Fig. 16. (ii) The redox ratio of scytonemin, ReScy/OxScy, of these filaments classified as [w/ OxDS, +UVA] were substantially lower than those of filaments classified as [w/ ReDS, no UVA] shown in Supplementary Fig. 11. Although the inclusion of the ReScy in the spectral decomposition of the filaments treated with UVA was positively supported neither by the SVD spectra (as in Extended Fig. 4) nor by the raw cellular spectra (as Fig. 2 in the main text), ReScy was included in the spectral decomposition, and the resultant ReScy distribution was plotted here for a fair comparison of the filaments between the different filament types.

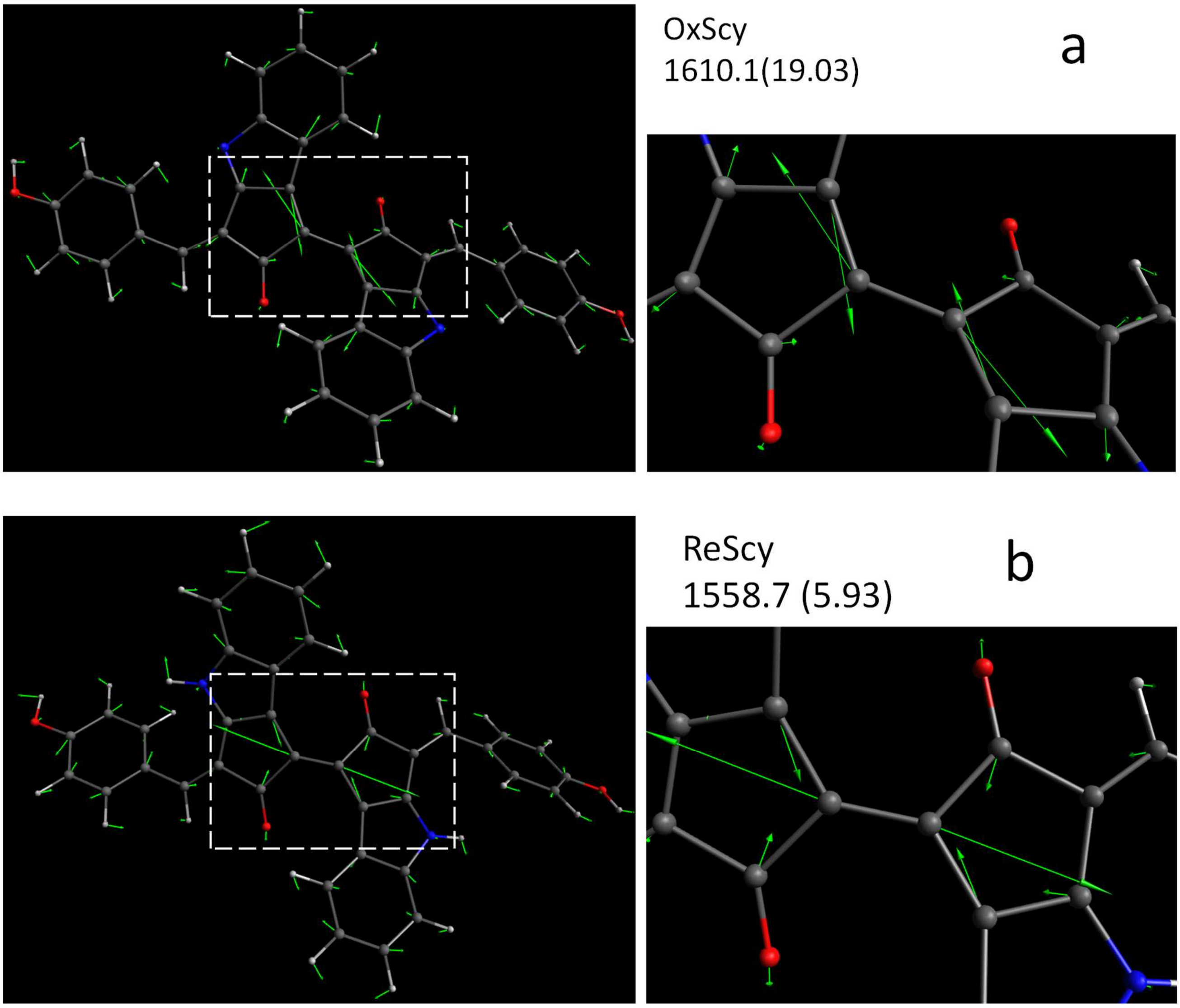


**Supplementary Fig. 13. Theoretical visualization of one vibrational mode that is mainly associated with the carbon-carbon bond (C1-C22) of the oxidized or reduced scytonemin** (See Extended data Fig. 1 for the atomic numbering). (a) The mode in the oxidized scytonemin (OxScy) with a Raman shift of 1610 $cm^{-1}$. (b) The mode in the reduced scytonemin (ReScy) with a Raman shift of 1559 $cm^{-1}$. The green arrows show exaggerated atomic displacements by the vibration. Left panels show the whole molecules, in which the regions enclosed by broken-line rectangles are magnified as the right panels. The numbers in the brackets show the theoretically estimated Raman scattering intensities following the same definition as in the Extended data Fig. 8.

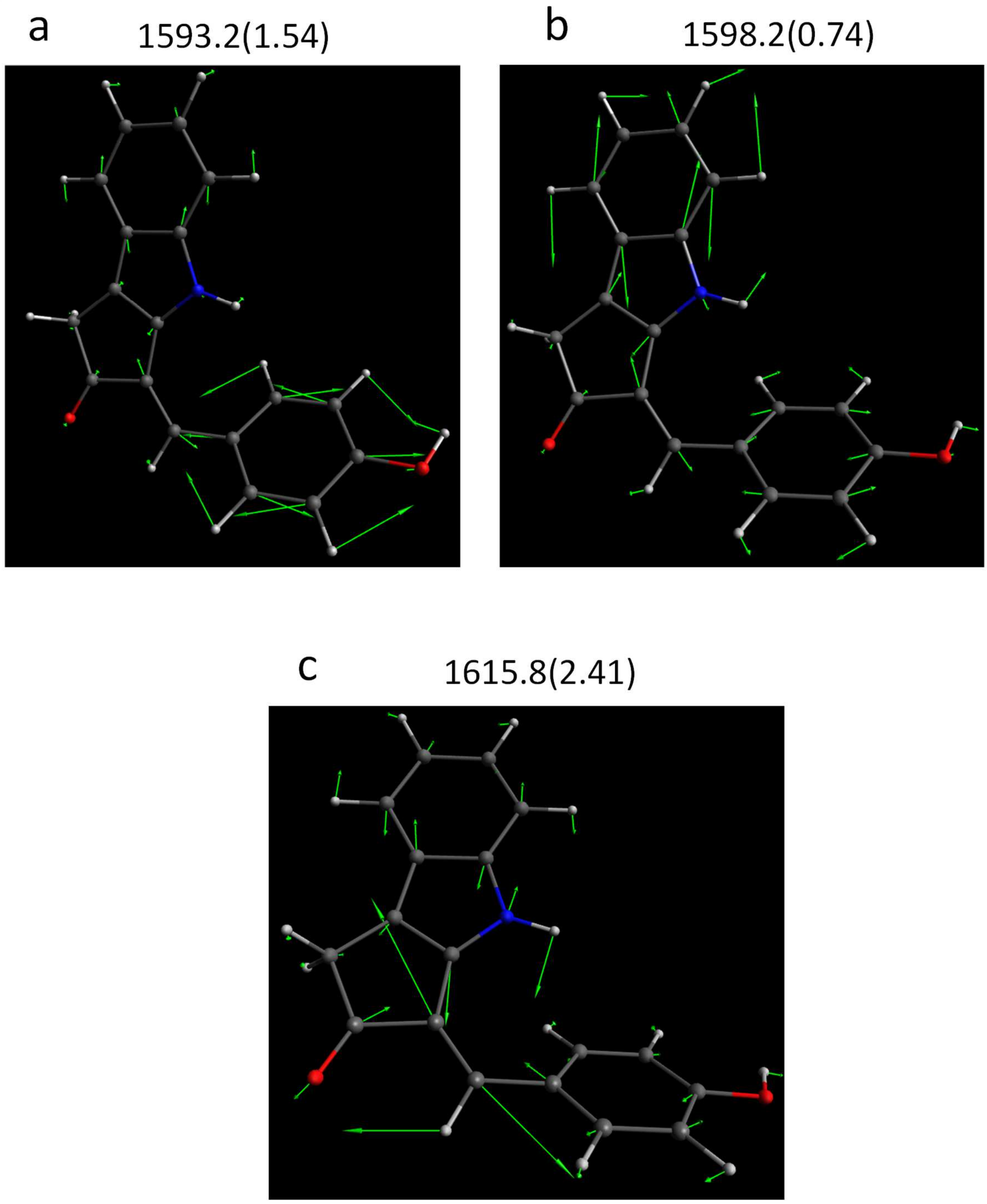


**Supplementary Fig. 14. Theoretical visualization of three vibrational modes of the monomeric scytonemin (MmScy)**. (a) and (b): These modes with Raman shifts of 1593 cm$^{-1}$ and 1598 cm$^{-1}$, respectively, are mainly attributed to C-C and/or C=C stretching vibration modes coupled with locally in-plane C-H bending that are mainly associated with the two aromatic rings. (c) This mode is mainly attributable to the C=C stretching vibration of the vinyl bond, corresponding to C4=C13 or C36=C45 in the dimeric scytonemins (Extended data Fig. 1). The green arrows show exaggerated atomic displacements by the vibrations. The numbers in the brackets show the theoretically estimated Raman scattering intensities following the same definition as in the Extended data Fig. 8. It is to be noted that we found three corresponding vibrational modes with similar distortions in the OxScy that have very similar frequencies.

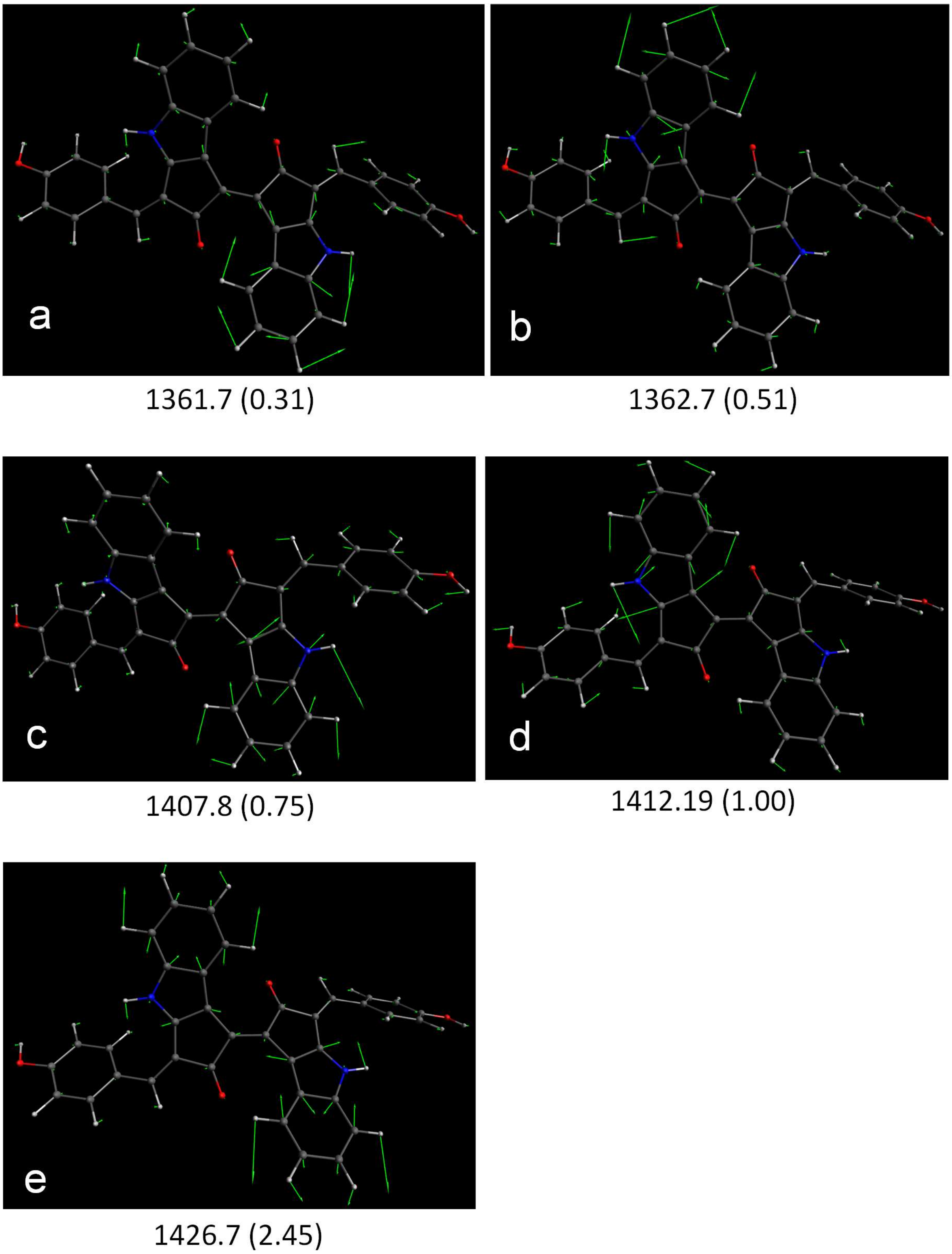


**Supplementary Fig. 15**

Theoretical visualization of five vibrational modes of the reduced scytonemin (ReScy) that are candidates for the experimentally observed Raman band at 1365 – 1366 cm$^{-1}$ in the *Rivularia* filament and in the solid-state. The green arrows show exaggerated atomic displacements by the vibration. Predicted Raman shifts in cm$^{-1}$ are shown below the images. The numbers in the brackets show the theoretically estimated Raman scattering intensities following the same definition as in the Extended data Fig. 8. The Raman shifts in (a) and (b) show the best match to the experimentally observed ones, but the Raman intensities are weaker than those in (c) – (e). All five vibrations included more or less locally in-plane N-H bending motion (at the atomic position of no. 6 or 38 in Extended data Fig. 1) that are sensitive to the redox change between ReScy and OxScy.

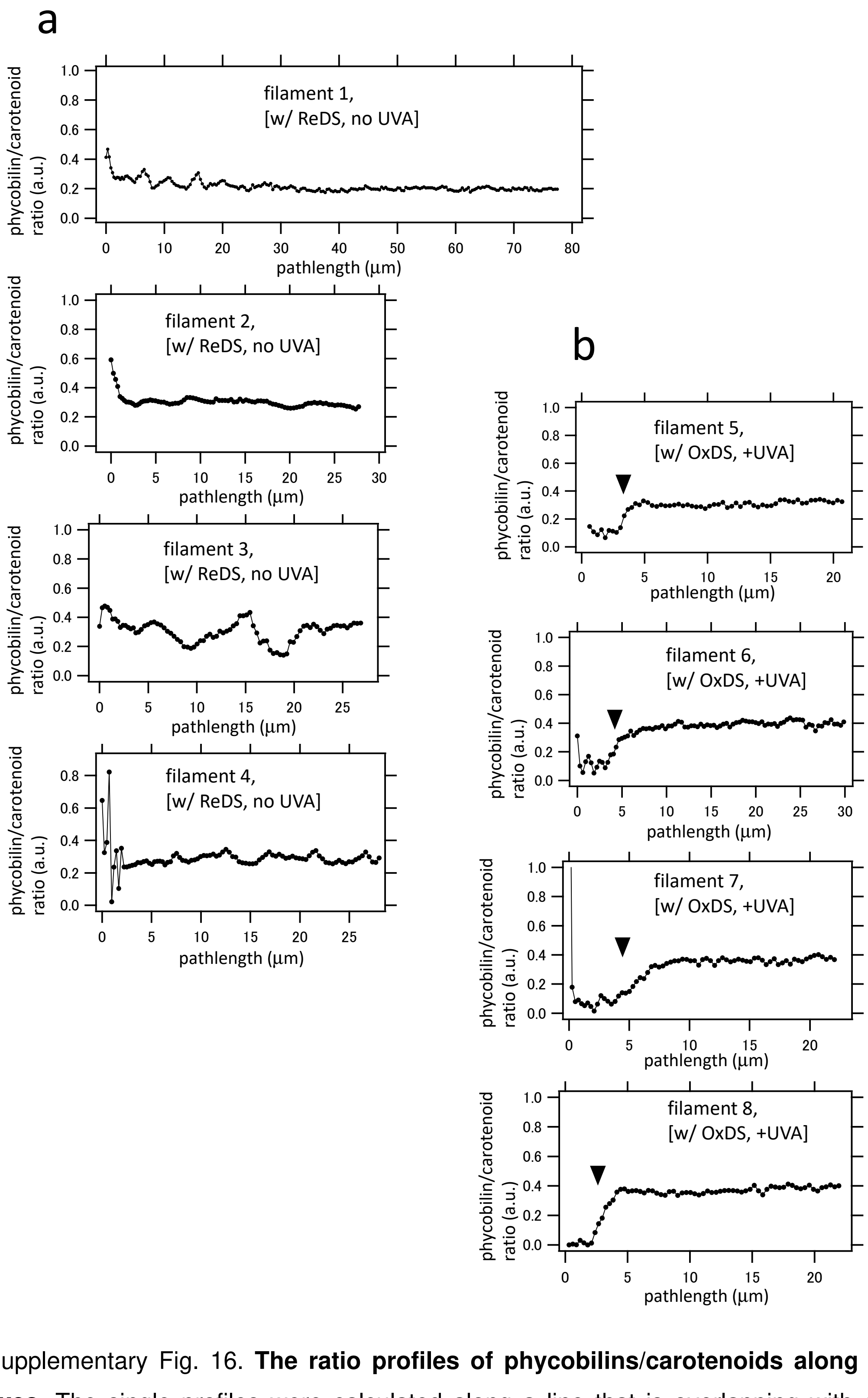


Supplementary Fig. 16. **The ratio profiles of phycobilins/carotenoids along longitudinal filament axes.** The single profiles were calculated along a line that is overlapping with centers of cytoplasmic regions and cell junctions (See Fig. 1g for examples). (a) Four filaments classified as [w/ ReDS, no UVA] (designated by filament numbers from 1 to 4) were selected. The filament numbers are corresponding to those in Supplementary Fig. 11. (b) Four filaments classified as [w/ OxDS, +UVA] (designated by the filament numbers from 5 to 8) were selected. The filament numbers are corresponding to those in Supplementary Fig. 12. The position of heterocyst-vegetative cell junction is indicated by black downward arrow heads. Clear decreases in the ratio of phycobilins/carotenoids around the terminal region (heterocyst position, about 0 – 4 μm) in comparison with the region of vegetative cells (>5 μm) were observed only in (b), but not in (a). Most of dark-sheathed filaments classified as [w/ OxDS, no UVA] also exhibited the decrease of phycobilins/carotenoids in the terminal region (heterocyst position). See also Table 1 in the main text.

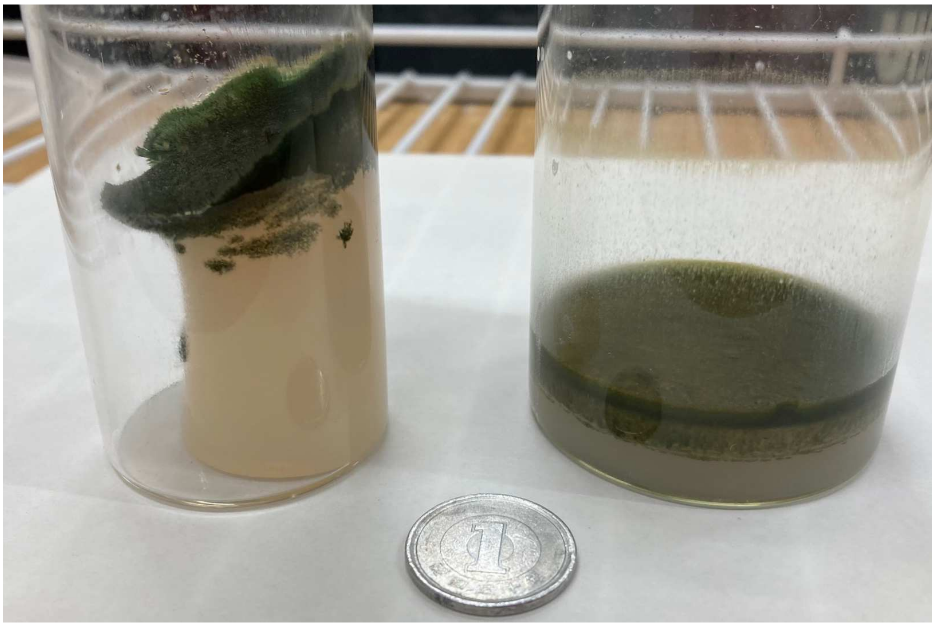

**Supplementary Fig. 17. Picture of thick accumulations of *Rivularia* filaments on agar media in glass bottles.** The left bottle contains *Rivularia* filaments on an agar layer based on $BG11_0$, and the right bottle contains *Rivularia* filaments on an agar layer based on BG11 containing fixed nitrogens. The Japanese coin put near the bottles has a diameter of 20 mm. It is to be noted that we have never observed such a thick accumulation of filaments in the cases of *Anabaena variabilis*, *Anabaena* sp. PCC 7120, and *Nostoc punctiforme*, when they are grown under very similar conditions. The matured filaments of these three species lack obvious polarity and contain multiple heterocysts with semiregular intervals in a single filament. This comparison suggests that *Rivularia* M-261 filaments tend to form a 3D colony, although we cannot yet describe the microscopic morphology of the colony in details.

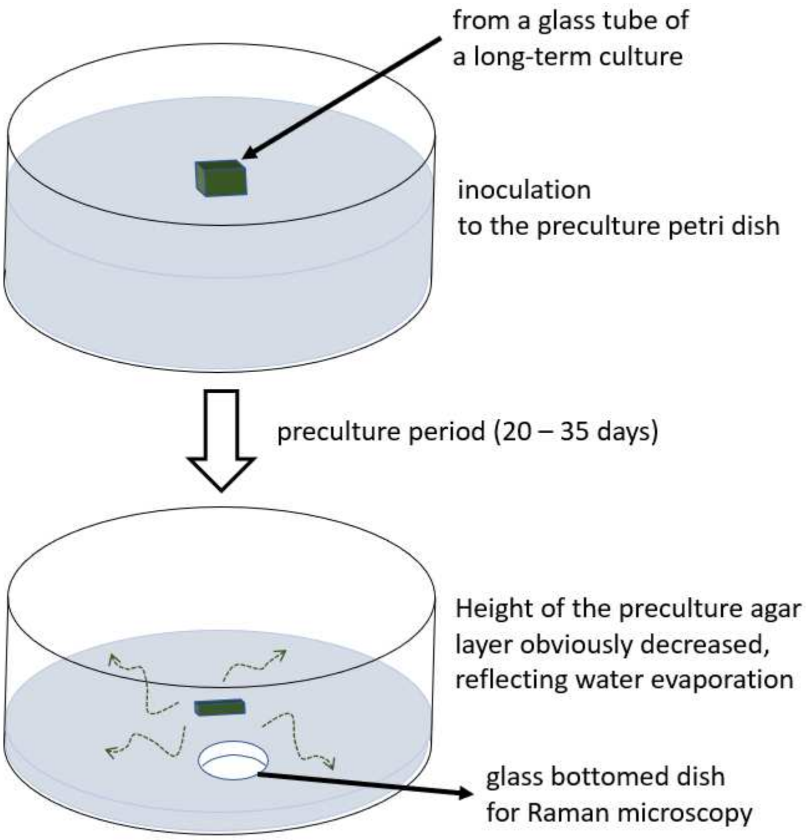


**Supplementary Fig. 18. Schematic drawing of a preculture of *Rivularia* filaments before microscopic observations.** At the inoculation, an agar block was transferred from a long-term culture in which *Rivularia* filaments were grown to a high cell density (e.g., Supplementary Fig. 17) under exogenous-nitrogen-depleted conditions ($NG11_0$ medium). The preculture polystyrene petri dish with an internal diameter and depth of about 54 and 12 mm, respectively, also contained the $BG11_0$ medium having water content in mass of higher than 98 % at the beginning, but the height of the agar layer reflecting the water content obviously decreased after the preculture period of 20-35 days. This means that there were desiccation and/or nutritional stresses to the filaments classified as [w/ OxDS, no UVA] and [w/ OxDS, no UVA], while there was no UVA illumination. Upon the inoculation to the new agar in the preculture dish, highly motile filaments (hormogonia) were differentiated from mature *Rivularia* filaments on the inoculated old small agar block, and they move to intact agar surface far from the mother filaments. The agar blocks for microscopic observations were cut off from regions away from the originally inoculated old block by an approximate distance of 15 – 30 mm. See also Method section in the main text.

References for the supplementary materials.


1. Tamamizu, K. & Kumazaki, S. Spectral microscopic imaging of heterocysts and vegetative cells in two filamentous cyanobacteria based on spontaneous Raman scattering and photoluminescence by 976 nm excitation. *Biochimica et Biophysica Acta (BBA)-Bioenergetics* **1860**, 78-88 (2019).
2. Merlin, J.C. RESONANCE RAMAN-SPECTROSCOPY OF CAROTENOIDS AND CAROTENOID-CONTAINING SYSTEMS. *Pure and Applied Chemistry* **57**, 785-792 (1985).
3. Nozue, S., Katayama, M., Terazima, M. & Kumazaki, S. Comparative study of thylakoid membranes in terminal heterocysts and vegetative cells from two cyanobacteria, Rivularia M-261 and Anabaena variabilis, by fluorescence and absorption spectral microscopy. *Biochimica et Biophysica Acta (BBA)-Bioenergetics* **1858**, 742-749 (2017).
4. Paulo, C. & Dittrich, M. 2D Raman spectroscopy study of dolomite and cyanobacterial extracellular polymeric substances from Khor Al-Adaid sabkha (Qatar). *Journal of Raman Spectroscopy* **44**, 1563-1569 (2013).
